# Ecology and demographic structure of an extinct ibex population in Upper Palaeolithic Italian Alps

**DOI:** 10.1101/2025.04.01.646553

**Authors:** Elena Armaroli, Francesco Fontani, Rocco Iacovera, Elisabetta Cilli, Adriana Latorre, Donata Luiselli, Sara Silvestrini, Gabriele Terlato, Giampaolo Dalmeri, Alex Fontana, Nicola Nannini, Hubert Vonhof, Lucio Calcagnile, Gianluca Quarta, Rossella Duches, Eugenio Bortolini, Anna Cipriani, Stefano Benazzi, Federico Lugli, Matteo Romandini

## Abstract

Alpine Upper Palaeolithic contexts exhibit specialised subsistence strategies, heavily dependent on *Capra ibex*. Among them, the rock shelter Riparo Dalmeri stands out, with *C. Ibex* dominating faunal remains across all occupation phases, spanning the Pleistocene/Holocene transition. This evidence positions Riparo Dalmeri as a key site for exploring the interdependence between human groups and *C. ibex* during one of the most critical climatic and cultural shifts in human evolution. Here, we present the first multidisciplinary study on Late Palaeolithic *C. ibex* teeth from Riparo Dalmeri, integrating direct radiocarbon dating, isotope (^87^Sr/^86^Sr, δ^13^C, δ^18^O), proteomic, and aDNA analyses. We generated the earliest aDNA sequences for *C. ibex* and contextual evidence on mobility, seasonality, and sex ratios. We found that most *C. ibex* were local to the area despite consistent human presence. They reveal significant dietary differences between sexes as well as increased seasonality at the Pleistocene-Holocene transition. Our results identify Riparo Dalmeri as an extinct branch of the ibex mtDNA phylogeny, offering unprecedented insights into ibex ecology and evolution that resonate with present-day issues on the conservation of this species in the face of climate change.

## Introduction

The Alpine ibex (*Capra ibex*) is undoubtedly a symbol of the Alps, with its presence attested since the Palaeolithic^1^. Nevertheless, human activities have long posed significant threats to its survival. Overhunting drove the Alpine ibex to near extinction in the early 19th century, with repopulation only occurring recently from a small population that survived in the Gran Paradiso National Park (Italy)^2^. While the Alpine ibex is no longer considered an endangered species^3^, its natural habitat remains at risk, threatened by global warming^4,5^. Recent DNA studies highlighted a complex population history for Alpine ibex in Europe, identifying major bottleneck events resulting from both environmental fluctuations and excessive human exploitation ^6,7^. These results importantly demonstrate the value of ancient *C. ibex* genomic diversity and population size as proxies for reconstructing past ecological changes. In the Italian Alps, *C. ibex* played a key role in human subsistence since the Upper Palaeolithic, as testified by the widespread presence of remains in both valley floor and mid-altitude sites^8–10^.

Among these sites, Riparo Dalmeri stands out as an exceptional case study. Located in the Northeastern Italian Alps (Trentino region) at 1240 m a.s.l., it was repeatedly occupied by Late Epigravettian hunter-gatherers focused on ibex hunting^11^. Radiocarbon dating of charcoal and bone samples has identified three main phases of human occupation, spanning the critical climatic transition between the Pleistocene and the Holocene (13400-11500 cal. BP^12^). Despite detailed studies of these three phases, significant uncertainties remain, particularly regarding human presence in the rockshelter during the Younger Dryas cold event (ca. 12900- 11500 cal. BP^13^). These uncertainties are primarily due to the complex stratigraphy of the site^12,14^ (Supplementary Note 1). Riparo Dalmeri can be defined as a specialised ibex-hunting site^15^, with *C. ibex* remains representing 80% to 93% of identified species across all three phases of occupation (Supplementary Note 2). This pattern is in sharp contrast with other coeval sites in northern Italy^8^, making Riparo Dalmeri a key site for understanding human-ibex relationships during a period of climatic transition.

Here, we address Palaeolithic *C. ibex* ecology and evolution through a multidisciplinary study on ibex teeth recovered within the entire stratigraphic sequence of Riparo Dalmeri. Six new direct radiocarbon dates were obtained to refine our understanding of human occupation of the site at the end of the Upper Palaeolithic. Strontium (^87^Sr/^86^Sr), carbon (δ^13^C) and oxygen (δ^18^O) isotope analyses are used to provide a detailed picture of *C. ibex* diet and mobility, and to reconstruct the paleoclimate of this transitional period in the Italian Alps. Ancient DNA data - coupled with proteomic analysis of tooth enamel - are used to determine the sex of individuals. We also explore the genetic population structure of Riparo Dalmeri ibex and reconstruct its mitogenomic phylogeny. Our findings are discussed considering 1) the complex relationship between the hunter-gatherers of Riparo Dalmeri and their favourite prey and 2) the implications of this multi-proxy study for understanding the challenges faced by Alpine ibex in the current period of climate change.

## Results

### Revisiting the stratigraphic succession at Riparo Dalmeri

The zooarchaeological material analysed in this study was selected to secure an even representation of the three identified phases of human occupation at the site (Extended Table 1). New radiocarbon dates (Supplementary Table S1) confirm higher intensity of human activities during Phase 1 (13550-12950 cal. BP, 1*σ*) and Phase 2 (12950-12600 cal. BP, 1*σ*), i.e., the Late Glacial interstadial. Trampling and other post-depositional processes may have caused some vertical and horizontal displacement of anthropogenic material, reflected in occasional mismatches between calibrated and stratigraphically modelled temporal intervals (Extended Figure 1). However, the overall integrity of this stratigraphic section is supported by field observations, lithic refittings, and the coherent spatial distribution of findings. The new dates from the upper layers of the third phase (13100-11450 cal. BP, 1*σ*) partially overlap with the previous phases, suggesting a more complex taphonomic history of the external portion of the stratigraphy, despite its integrity and coherence with the Pleistocene/Holocene transition and the Younger Dryas.

### Paleoclimate reconstruction and the alpine ibex ecology

The complete list of isotopic results is available in the Supplementary Information (Supplementary Table S2). The local ^87^Sr/^86^Sr baseline ranges between 0.70798 and 0.70834 (mean = 0.70818, SD = 0.00013; n = 5). Tooth enamel ^87^Sr/^86^Sr values of the whole dataset span between 0.70794 and 0.70886 (mean = 0.70818, SD = 0.00021; n = 28) (Figure 1a), with no significant variations across different age classes, sexes, or occupation phases of the site. Ibex teeth from the first phase display values between 0.70825 and 0.70796, with a mean of 0.70812 (SD = 0.00012; n = 4). ^87^Sr/^86^Sr values from the second occupation phase range between 0.70886 and 0.70794, with a mean of 0.70818 (SD = 0.00023; n = 16, including four different tooth classes from the same individual). Samples from the third phase of occupation yielded a mean of 0.70816 (SD = 0.00021; n = 9), ranging between 0.70859 and 0.70794. Only the two most radiogenic samples, RD_8203 (^87^Sr/^86^Sr = 0.70886; second phase) and RD_8173 (^87^Sr/^86^Sr = 0.70859; third phase), are considered outliers. Yet, the majority (twenty-six out of twenty-eight) of ^87^Sr/^86^Sr values fall within the local range defined by the baseline samples and the Italian isoscape^16^. Thus, ibex teeth from Riparo Dalmeri can be considered a reliable paleoclimatic proxy for studying the local environment through δ^13^C and δ^18^O isotope analysis of hydroxyapatite carbonate-moiety.

**Figure 1.**
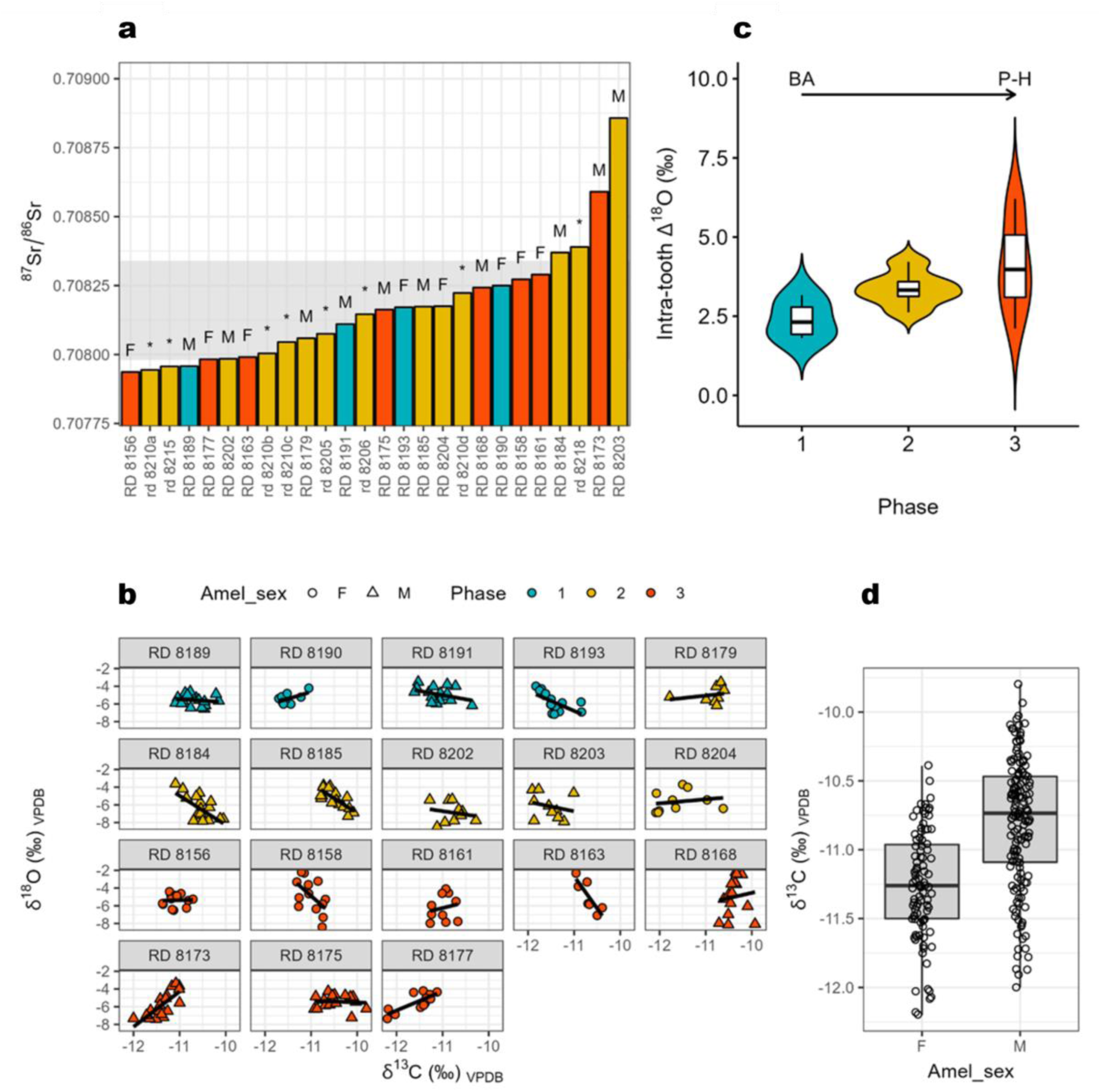
**a)** ^87^Sr/^86^Sr ratios for each individual (sex is labelled as “M” = male and “F” = female; “*” = deciduous tooth); the grey area represents the measured local range (obtained from 3 micromammal teeth, 1 hare bone and 1 ibex dental root), yet the Italian isoscape indicates a much larger range for the local Sr isotope signature at Dalmeri, possibly including the whole dataset of *C. ibex*; **b)** Intra-tooth δ^18^O vs δ^13^C values for each individual, coloured by phase (light blue = 1, yellow = 2, orange = 3), with symbols representing sexes (circles = female, triangle = male); **c)** Calculated intra-tooth Δ^18^O variability (max-min) for each individual, clustered by phase of site occupation, between the Bølling–Allerød (BA) interstadial and Pleistocene-Holocene (P-H) transitional period; d) δ^13^C values by ibex sex (Wilcoxon rank-sum test *p* < 0.01).

Sequential C and O isotope analysis of eighteen selected ibex teeth (n = 243 data points) yielded δ^13^C_VPDB_ values ranging from -12.2 to -9.8‰ (mean = -10.9‰), with no significant intra- tooth variation (∼1‰; Figure 1b; Extended Figure 2). No remarkable differences were observed among occupation phases (mean = -11.1, -10.8 and -10.9‰, respectively; Figure 1c). A significant difference (Wilcoxon rank-sum test *p* < 0.01) between females and males suggests dietary differences or habitat related differences (Figure 1d). Overall, these δ^13^C values are typical of C_3_ plant feeders^17^, consistent with Western European vegetation patterns^18^. The δ^18^O_VPDB_ values range between -8.5‰ and -2.2 (mean = -6‰) with high intra- tooth variation (up to 6.2‰, sample RD_8158; Figure 1b; Extended Figure 2), suggesting seasonal fluctuations in the local environment^19^. No significant difference between sexes was observed (Wilcoxon rank-sum test *p* = 0.42).

Overall, the variability observed in enamel δ^18^O data (recalculated as ingested water) match the modern predicted monthly precipitation δ^18^O time-series from Piso.Ai^20^ (Extended Figure 3) at the coordinates of Riparo Dalmeri. Given the known ibex tooth formation time^21^ and the observed seasonal δ^18^O peaks, individual teeth likely capture between six to twelve months of life history. Interestingly, while mean δ^18^O values are similar among the three occupation phases, the signal amplitude is significantly higher in the third phase. Air temperature estimates from oxygen isotopes suggest summer temperatures reach their highest (∼ 30°C) in summer and their lowest (∼ -5°C) in winter in the third phase. These estimates indicate colder conditions than today (Extended Figure 4). A statistically significant correlation between δ^18^O and δ^13^C values was observed in six out of eighteen samples (*p* < 0.05), with four showing negative correlations and two positive correlations (Extended Figure 5).

### The Dalmeri ibex mitochondrial phylogeny

Genetic data were generated from twelve well preserved teeth from Riparo Dalmeri (Supplementary Table S3). Additional samples were collected from the prehistoric sites of Riparo Cogola (NE Italy, n = 2) and Romagnano Loc III (NE Italy, n = 2) to provide a broader picture of genomic variability in Upper Palaeolithic Alpine ibex (Figure 2a).

**Figure 2.**
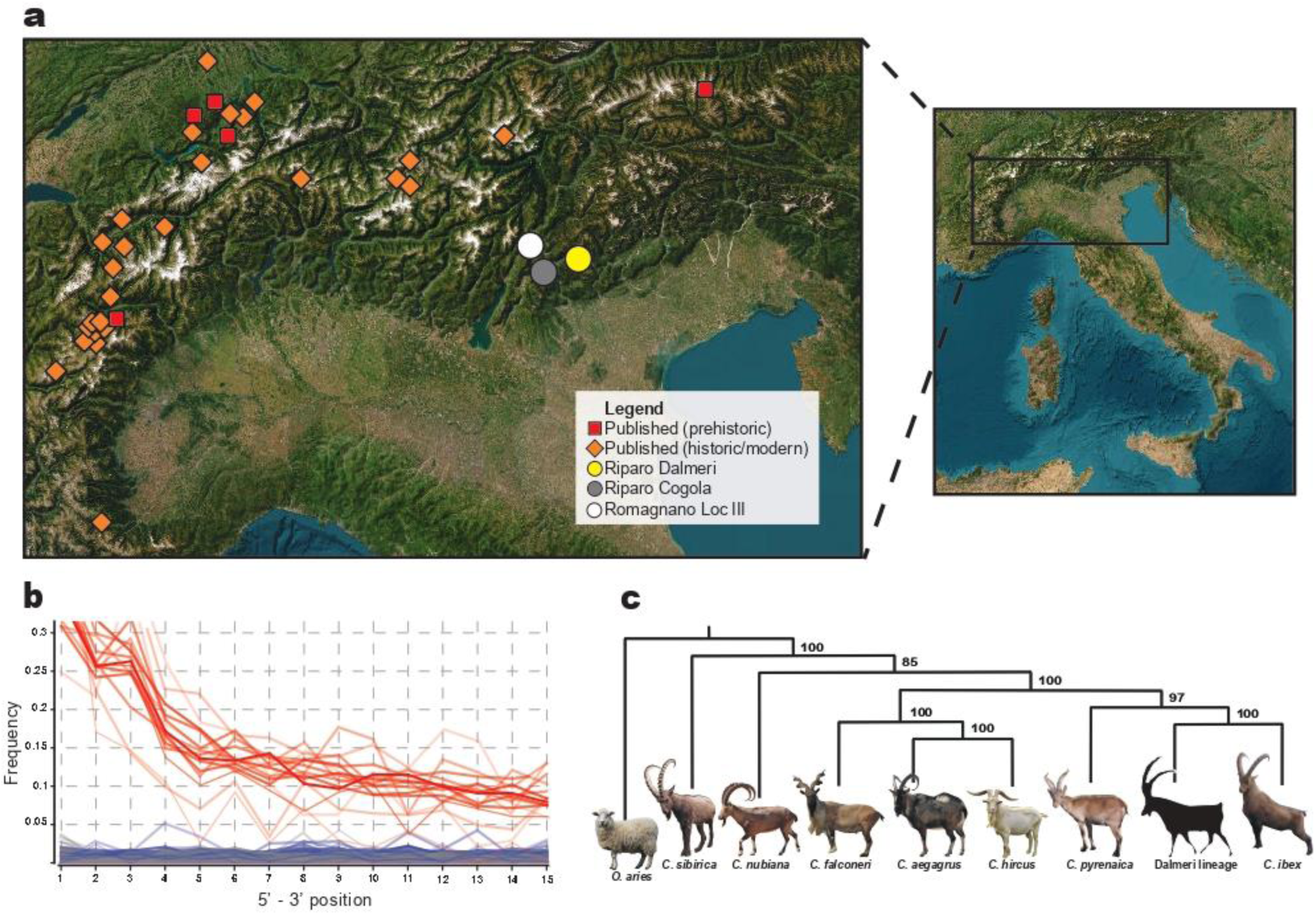
**a)** Geographic origin of the published and new ibex genetic data analysed in this study. Esri, World Topographic Map. Map was edited in Adobe Illustrator CC 2020; **b)** Recurrence of damage patterns in the form of C > T transitions (red lines) at 5’ end positions of single stranded DNA sequences for newly generated mitogenomes. Blurred lines are used for samples with mean coverage < 5x; **c)** A simplified Maximum Likelihood tree summarises the phylogenetic history of *Capra* species, with samples from Riparo Dalmeri clustering in the branch of *Capra ibex*. *Ovis aries* is used as the outgroup.

The samples reported 0.213% to 43.06% of preserved endogenous DNA when mapped to the nuclear reference genome of *Capra hircus* (ARS1.2). For each individual the total number of unique reads mapped on the Alpine ibex reference mitogenome (NC_020623.1) was consistently greater compared to mapping statistics on the mitochondrial references of other species of the *Capra* genus. Given the low coverage nature of the data (from 0.2x to 0.0003x), nuclear genomes were only used to assess the sex of the individuals. Analysis of degradation confirms authenticity of aDNA (Figure 2b). We reconstructed mitogenomic information from eleven of the twelve samples of Riparo Dalmeri, with mean coverages ranging 2x to 40x. Only samples with mean coverage ≥ 10x were retained for phylogenetic analysis (n = 6), so all samples from Riparo Cogola and Romagnano Loc III were discarded (Supplementary Note 3).

The Maximum Likelihood (ML) tree computed from a dataset of modern and ancient *Capra* species (“Capra_all”) shows that *C. ibex* and *C. pyrenaica* cluster together, forming a well-supported monophyletic clade for the European wild ibex (Extended Figure 6) that is clearly distinct from the remaining goat species from Eurasia and North East Africa, as well as from the domestic goat (*C. hircus*). Within this clade Riparo Dalmeri forms a sub-clade with sister group *C. pyrenaica*, while all historic *C. ibex* samples are located between Riparo Dalmeri and modern samples.

The Bayesian phylogeny (Figure 3a) supports the same tip topology and the same monophyletic origin of the Pyrenean specimens, with estimated divergence time between *C. ibex* and *C. pyrenaica* at 43500-31600 years BP (95% Highest Posterior Density, HPD). All modern *C. ibex* samples cluster together (*p* = 0.91), including the historic Swiss (Zu01 and Be4) and Italian Gran Paradiso samples. Riparo Dalmeri segregates from other ancient samples including the geographically closest Ötzi_stomach (*P* = 0.73), with an estimated divergence time of 41100-29800 years BP (95% HPD).

**Figure 3.**
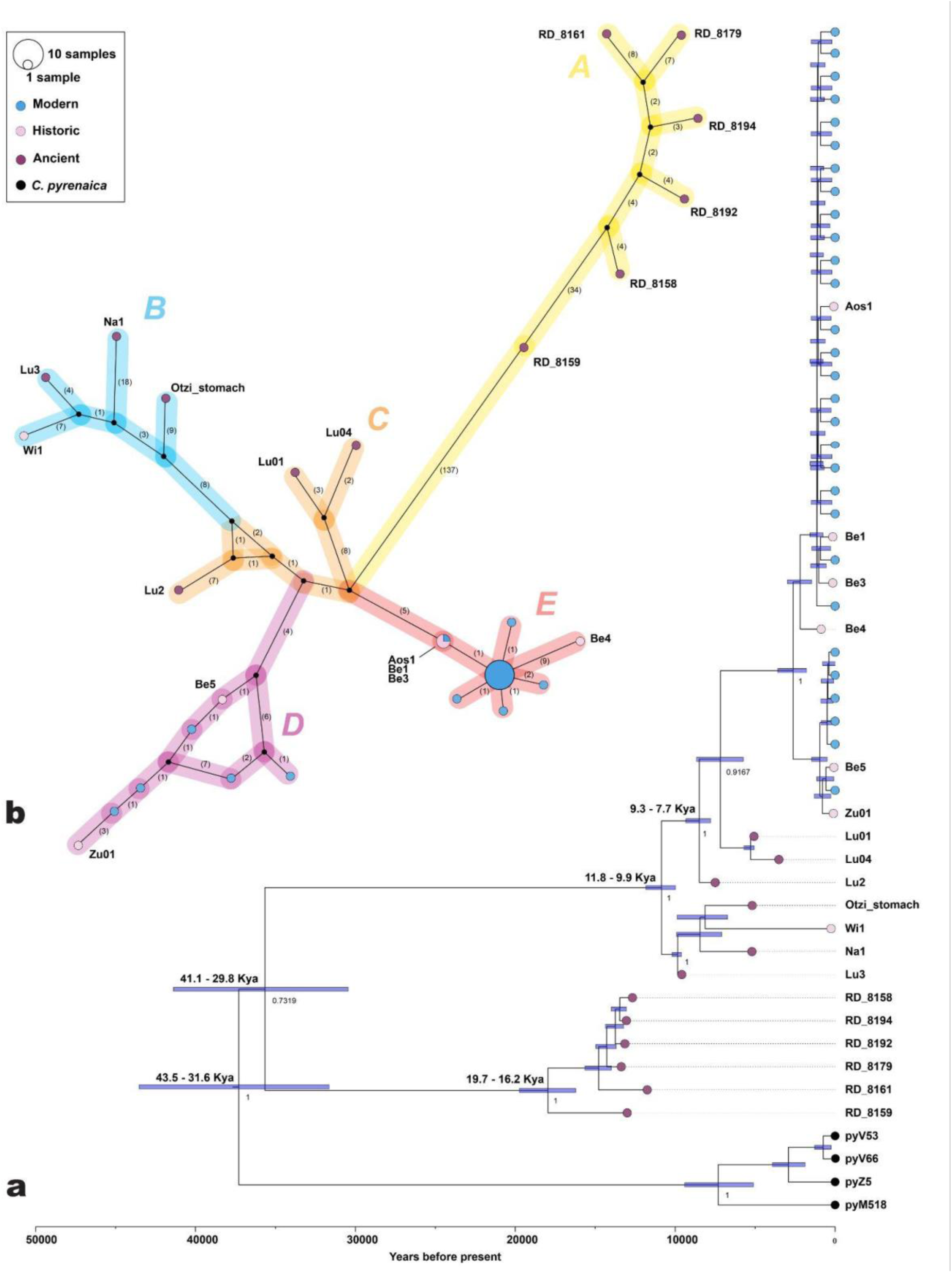
**a)** The Bayesian tree, constructed using the “Capra_ibex/pyrenaica” dataset. Branch labels denote the posterior probabilities and node labels indicate inferred split times. All key parameters exhibited effective sample sizes (ESS) exceeding 200, indicating that the MCMC chains were well-mixed and stabilised. The light blue blocks surrounding the nodes represent the 95% HPD intervals. Branch tips for *Capra ibex* are color-coded: blue for modern samples, pink for historical samples, and purple for ancient samples. Black branch tips represent samples of *Capra pyrenaica*. **b)** Haplotype network of the mtDNA sequences from the dataset “Capra_ibex”. A single substitution step is represented by a single number connecting two haplotypes. Non-sampled intermediary haplotypes are represented by black vertices connecting two or more haplotypes. The chronological context of the samples can be identified by their colours, as indicated in the legend.

The same topology is further supported by our mtDNA network (Figure 3b), which displays a total of twenty-six distinct haplotypes divided into five haplogroups. Among them, Haplogroup A only contains the ancient samples from Riparo Dalmeri, and the other ancient samples are segregated from modern and historic samples into distinct haplogroups (Supplementary Note 3). The Bayesian skyline shows that the population size did not experience significant fluctuations after the end of the Last Glacial Maximum (Extended Figure 7). Conversely, a visible population decline coincides with the large-scale growth of human populations in historic times. Notably, these results are consistent with the pairwise comparison analysis; the average genetic distances within Riparo Dalmeri (haplogroup A = 0.135) and the other ancient samples (haplogroups B = 0.127, and haplogroup C = 0.091) are approximately three times greater than that of the modern and historical samples in haplogroups D (0.044) and E (0.014). However, the average genetic distances in haplogroup A compared to haplogroups B and C are only 0.008% and 0.043% greater, respectively. Pairwise distances among ancient samples (Extended Table 2) show consistently higher dissimilarity between Riparo Dalmeri and other ancient samples.

### A combined approach for sex estimate in prehistoric Alpine ibex

Proteomic sex estimation was performed exploiting the sex-specificity of AMELX and AMELY protein isoforms encapsulated in tooth enamel. The presence of both unique AMELX and AMELY peptides in fifteen fossil samples indicates male individuals while the remaining eight samples, which only contained AMELX peptides, are likely female. The detailed procedure is described in the Supplementary Text (Supplementary Note 4).

Sex was also identified on ten of the eleven specimens sequenced for ancient DNA. Even at extremely low coverage (mean < 0.0003), DNA-based determination with both the Rx and Ry values matched five out of six results obtained by amelogenin analysis on overlapping samples (Supplementary Note 5). Overall, genetic results unambiguously identified six male and four female individuals. Due to low coverage, RD_8158 reported borderline results on both Rx (0.734, CI 0.687-0.782) and Ry (0.026, CI 0.023-0.029) estimates. However, amelogenin analysis, along with an extremely low quantity of DNA sequences mapping to Y-chromosome regions (n = 313), suggests the individual was female. Read depth statistics for Y-chromosome, X-chromosome and autosomal regions further support the consistency of the methods (Supplementary Table S4).

## Discussion

Previous research has suggested that Riparo Dalmeri was a seasonal site specialised in ibex hunting^22^, a camp site where all family members – including children^23,24^ – lived seasonally basing most of their subsistence on ibex^15,25^. The impressive abundance of *C. ibex* remains (see Supplementary Note 2) and the recurrent presence or representation of this species in dwelling structures, hearths, ornaments, and portable art, make Riparo Dalmeri a unique case study in the entire European continent for studying the interaction between this taxon and humans on the verge of a pivotal climatic and cultural tipping point.

In this work new, direct radiocarbon dates, together with literature data, confirm that Riparo Dalmeri was frequented during the Late Glacial, primarily during the Bølling-Allerød interstadial (13550-12950 cal. BP, 1*σ*; first and second phases), when Late Epigravettian hunter-gatherers began to recolonise the mountains. Despite the presence of reworked materials, the new dates (12950-11450 cal. BP, 1*σ*; second and third phases) suggest anthropic frequentation, and thus ibex hunting, also during the Younger Dryas or Pleistocene-Holocene transition. This is supported by the presence of lithic tools coherent with this chronological period (Supplementary Table S1; Supplementary Note 1).

We also explored the genetic footprint of Riparo Dalmeri ibex to provide new understanding of its population structure in light of the extensive pressure posed by specialised hunting activities. Our findings confirm previous evidence on the mitochondrial phylogenetic relationships among different *Capra* species. Here, *Capra ibex* forms a clearly distinct monophyletic clade. Based on radiocarbon dates and Bayesian phylogeny, we now estimate that the divergence between *Capra ibex* and *Capra pyrenaica* occurred ∼37550 years BP, post-dating previous estimates by nearly 10000 years. In the Bayesian tree, the ibex from Riparo Dalmeri form a clearly distinct branch, possibly representing an ancestral, geographically isolated, and extinct population within the ibex mtDNA phylogeny. This structure may have consolidated during the Last Glacial Maximum, when ice coverage reached its peak, extending over large parts of Northern Europe and the Alps^26^. When compared against samples from the coeval and geographically close sites of Riparo Cogola and Romagnano Loc III, Riparo Dalmeri still emerges as a segregated population – although the low coverage of the former hampers statistical support and warrants caution when interpreting these results. On the other hand, the geographic and temporal distance of Riparo Dalmeri from the other ancient samples analysed in this study may explain at least part of their genetic segregation. This genetic model will benefit from the future inclusion of additional ancient sequences from both sides of the Alpine range.

As far as population size over time is concerned, Riparo Dalmeri shows levels of haplotype and nucleotide diversity consistent with other ancient samples, suggesting a stable density of *Capra ibex* during the Upper Paleolithic, despite intensive hunting and consistent human presence. Despite rise in temperatures and consequent changes in Alpine habitats, as well as increasing human activity^27^, our Bayesian analysis does not show a decrease in population size after the LGM. A drastic decline in intraspecific genetic diversity can only be detected between the 16th and 18th centuries due to increased anthropogenic pressure on the ecosystem, followed by a dramatic bottleneck which almost drove the species to extinction after the 18th century^7,28^.

Proteomic and isotope analyses carried out in this study shed light on the ecology of this extinct and isolated population in the broader context of the marked climatic variability characterising the end of the Late Glacial period. Here, for the first time, we compare amelogenin and aDNA results on sex determination in ancient animal samples. We identified eight females out of twenty-three samples, indicating that both sexes of different age classes were present around the site during human occupation of the shelter, likely coinciding with the ibex hunting season^22,29^. Ecological studies suggest that modern ibex primarily move seasonally through altitudinal shifts over relatively short distances (from 6 to 22 km)^30^. Even shorter distances have been observed in the autochthonous population of Gran Paradiso National Park, showing a degree of fidelity to seasonal home ranges^31^. The low variability of our ^87^Sr/^86^Sr values, which align with the local baseline, supports the idea of limited seasonal movements^32^. Considering the low degree of modern ibex mobility and the significant variability of ^87^Sr/^86^Sr values in the area^16^, the two samples identified as possible outliers (RD_8203; RD_8173) likely reflect short-range mobility. Notably, both outliers are males. Modern studies show that present-day male and female ibex select different habitats throughout the year, with males using larger home ranges^1,33,34^.

The relatively high δ^13^C values in the Dalmeri samples are typical of semi-open environments^35^ and are consistent with recently published δ^13^C values from ibex bone collagen (SU 26b-c)^36^. Small intra-tooth variations could result from the narrow isotopic range documented in C_3_- dominated environments of temperate and boreal ecosystems^37^, but might also reflect the limited mobility indicated by Sr isotopes. The lack of significant differences across the three occupation phases (mean values ∼11‰) may be related to these factors.

However, we cannot exclude that the ∼1‰ intra-tooth variation reflects altitudinal mobility. A decrease in ^13^C discrimination with altitude has been extensively documented in C_3_ plants^38,39^ and various animal species, suggesting potential altitudinal shifts of about 200 meters^37^. Altitudinal mobility is also often indicated by a negative correlation between δ^13^C and δ^18^O values in tissues of animals living in mountain environments^40,41^. However, in our dataset, 12 out of 18 ibex samples do not show a significant correlation between these two values (p > 0.05; Extended Figure 5). Sex-related differences in altitude use do not appear to explain these variations. Among the six samples that show a significant correlation (p < 0.05), both sexes are represented, which is consistent with previous studies suggesting no major sex differences in altitude use^32^. Conversely, a significant δ^13^C difference between sexes in fossil *C. ibex* individuals (Figure 1d) aligns with ecological data on modern ibex. The strong sexual dimorphism, along with other factors like pregnancy and the presence of ibex kid, leads to habitat segregation and distinct plant consumption patterns for much of the year^33,42^.

From an environmental perspective, oxygen isotopes provide key insights into the paleoclimate at Riparo Dalmeri. Because ibex are obligate drinkers^43^, their δ^18^O values mainly reflect local water sources, which are closely linked to local environmental parameters^44^. Lower values correspond to winter and higher values to summer months^19^. All ibex samples show a clear sinusoidal pattern in their δ^18^O values, indicating seasonal changes in the local environment, with no notable differences between males and females. Interestingly, while the highest intra-tooth δ^18^O values are comparable in the three occupation phases, the lowest δ^18^O values occur only in the third phase. As a result, most samples from the third phase display a larger amplitude (‰) of δ^18^O values. This increased amplitude suggests that the environment experienced greater seasonal changes during this period compared to the Bølling-Allerød interstadial. These findings seem to reflect the climatic shifts associated with the onset of the Younger Dryas and the subsequent Pleistocene-Holocene transition. Estimated air temperatures further support this interpretation. In the third phase, the highest temperature reached ∼30 °C and lowest temperatures dropped to ∼-5 °C, mirroring modern seasonal temperature fluctuations recorded at the meteorological station of Borgo Valsugana (∼400 m a.s.l., 25 km from Riparo Dalmeri; Extended Figure 4). We are aware that uncertainties in the model and the stratigraphy of the third occupation phase, especially for samples from the external succession, do not allow us to draw firm conclusions. However, several studies have reported that the Younger Dryas in the northern hemisphere was characterised by cool, dry winters and shorter summers than the Bølling-Allerød interstadial, even though summer temperatures where comparably high^45–47^. Our isotope data are consistent with this pattern.

These environmental and ecological findings complement our genetic insights. While our isotope data revealed increased seasonality in the environment during the third phase, likely linked to the onset of the Younger Dryas and the Pleistocene-Holocene transition, the genetic data suggest that such environmental pressures, along with intensified human hunting, might have driven shifts in ibex ecology. Changes in ibex “seasonality” and habitat use due to changes in climatic conditions, have been widely documented^1,31,32,34^. As the climate changed, the ibex population of Riparo Dalmeri likely adjusted their seasonal home ranges, which in turn could have influenced patterns of human activity at the site. Therefore, the archaeological evidence of sporadic human frequentations at the end of the Epigravettian could be related to a decreased availability of their favourite prey. Later, during the Holocene, ibex became restricted to high mountain refuges^48^, an occurrence echoed today in the Alpine region in response to heat stress^49^. The simultaneous action of anthropic pressure and the sudden change in the environment may have left the ibex population of Riparo Dalmeri insufficient time to adapt, leading to its extinction.

Together, the genetic, isotopic and archaeological evidence paints a coherent picture of how climatic shifts, human activities and geographical isolation shaped the evolutionary history and population dynamics of extinct *Capra ibex* at Riparo Dalmeri. We believe that this type of analysis should be more frequently integrated into archaeological and palaeoecological research, as it can help address critical issues such as the conservation and management of animal species threatened by human-induced climate change.

## Methods

### Material selection and archaeozoological analysis

A detailed description of the archaeological contexts considered in this study is provided in the Supplementary Note 1. Osteological samples from all the sites involved in the study were provided by the MUSE - Science Museum of the autonomous province of Trento (Italy). Both adults (n = 22) and young individuals’ teeth (n = 10) were selected. Since they are easier to recognize and more suitable for intra-tooth sequential sampling, only permanent third molars were selected for adult individuals. For young individuals, both permanent molars and deciduous premolars were selected. Age classes were determined following^25^. The determination of the minimum number of individuals (MNI) for each occupation phase was performed to avoid sampling teeth from the same individual. The MNI count was carried out considering both the side (i.e., left or right) and the tooth wear^21,50^. Moreover, the maximum anterior-posterior diameter (MAP) and the transversal diameter (TD) were measured on third molars only to allow comparison with the dimension of ibex teeth from other Italian Palaeolithic sites (Supplementary Tables 1 and 2).

To build the local ^87^Sr/^86^Sr baseline, plant and water samples could not be collected due to the current inaccessibility of the site. Therefore, we analysed archaeological fauna found during excavations at Riparo Dalmeri and provided by the MUSE (see^51^ for a review). These samples consisted of n = 1 *Lepus timidus* tooth enamel, n = 3 undetermined micromammal tooth enamel and n = 1 ibex tooth root. Detailed information about the samples is provided in the Extended Table 1 and Supplementary Note 2.

### Radiocarbon dating

Collagen was extracted from the samples by using the Longin protocol^52^ at the chemical laboratories of the Centre for Applied Physics, Dating and Diagnostics (CEDAD) Department of Mathematics and Physics “Ennio de Giorgi” University of Salento^53^. A fraction of the extracted collagen was separated from nine of the samples for the determination of the C:N ratio and for carbon and nitrogen stable isotope analyses which were performed by using an elemental analyser (EA-Mod. Flash 2000 HT by Thermo). The fraction selected for ^14^C analysis was combusted to CO_2_ in sealed quartz tubes with CuO and Silver wool and then reduced at 600°C to graphite with H_2_ on Fe powder used as catalyst. AMS ^14^C measurements were carried out with the 3 MV Tandetron at CEDAD-University of Salento (High Voltage Engineering Europa BV Mod. as in^54^. The measured ^14^C/^12^C isotopic ratios were corrected for isotopic fractionation by using the δ^13^C term measured on line with the accelerator, and for machine and chemical processing background. The conventional ^14^C ages were then calculated according to Stuiver and Polach^55^ and calibrated by using the last internationally accepted IntCal20^56^ calibration curve. Data (Supplementary Table S1) were analysed and phase duration was inferred through a contiguous stratigraphic Bayesian model for Phase 1 and 2 and an overlapping/independent sequence for the external Phase 3 using OxCal v4.4.4^57–59^ on the stratigraphic sequence published in^12^.

### Isotope analyses

Isotope analyses were carried out on dental enamel of 18 permanent ibex teeth from all three phases of occupation of the site, and 10 deciduous ibex teeth. ^87^Sr/^86^Sr isotope analysis was performed on both adult and young ibex individuals, as well as on 5 baseline samples. Sample preparation and Sr separation were carried out at the MeGic laboratory of the Department of Chemical and Geological Sciences of the University of Modena and Reggio Emilia. About 5 - 10 mg of each sample (‘bulk’) was dissolved in 3M HNO_3_. Strontium was separated from the matrix using chromatographic Teflon columns filled with 30 µl of Eichrom Sr spec resin^60^. The resin was cleaned with MilliQ water and conditioned using 3M HNO_3_ before sample loading. Cations (not Sr) were desorbed by percolating 3M HNO_3_. Sr was then eluted with several reservoirs of MilliQ water. The sample solutions obtained were diluted using 4% HNO_3_ and the ^87^Sr/^86^Sr isotopic compositions were measured using a Thermo Fisher Neptune MC-ICPMS, housed at the Centro Interdipartimentale Grandi Strumenti of the University of Modena and Reggio Emilia, as described in^61,62^. Mass bias normalization was performed through exponential law using an ^88^Sr/^86^Sr ratio of 8.375209^63^. Repeated analysis of NBS-SRM987 yielded an ^87^Sr/^86^Sr ratio of 0.710213 ± 0.000018 (2SD, n = 14). Samples were reported to an accepted NIST-SRM987 value of 0.710248^64^. Tooth enamel from the 18 adult specimens was sequentially sampled for oxygen (δ^18^O) and carbon (δ^13^C) isotope analysis. Samples ∼0.1 cm wide were drilled ∼0.2 cm apart starting from the occlusal surface (older enamel) to the enamel-root junction (younger enamel), overall corresponding to ∼1 year of life^21^. Depending on the tooth length and wear, we were able to obtain between 7 and 20 enamel samples per tooth. The analysis on the carbonate moiety of enamel hydroxyapatite was carried out at the inorganic stable isotope laboratory of the Department of Climate Geochemistry at the Max Planck Institute for Chemistry in Mainz. About 200-400 μg of enamel powder per sample was analysed on a Thermo Delta V mass spectrometer equipped with a GASBENCH-II preparation device. Within a run of 40 samples a total of 12 replicates of two in-house tooth enamel standards (AgLox, mammy) and one calcium carbonate (IAEA 603) were analysed. The standards yielded an external reproducibility (1SD) better than 0.15‰ for δ^18^O and 0.1‰ for δ^13^C. Standard material weights were chosen so that they span the entire range of sample weights. All data were reported to the VPDB (Vienna Pee Dee Belemnite) scale^65^.

### Inverse modeling and paleotemperature reconstruction

Due to the long time required for tooth mineralization and the sampling procedures, time-averaging and amplitude damping effects occur between the measured intra-tooth isotope values and the actual isotopic variations recorded in the enamel during tooth formation. Therefore, before using the intra-tooth δ^18^O values for seasonal paleotemperature estimation, inverse modelling was applied to recover the environmental input signal^66,67^. We are aware that this model was developed for continuously growing teeth, yet no specific model exists for med-sized bovids such as ibex (see^68^ and cfr.^69,70^).

The initial mineral content of enamel was set at 25%, enamel appositional length at 10 mm, and maturation length at 13 mm (see^71^). During the modelling workflow, a damping factor describing the damping of the isotopic profile amplitude needs to be iteratively chosen using an adjustment of the measured error term (E_meas_) to the prediction error (E_pred_). The adjusted damping factors here used ranged between 0.01 and 0.03. Carbonate δ^18^O values were converted to phosphate (δ^18^O_ph_)^72^ and then to δ^18^O of ingested water (δ^18^O_w_) using the formula for *Capra sp.* from^73^. The modelled input δ^18^O_w_ profiles are shown in Extended Figure 8. Summer, winter and mean air temperature estimation were obtained for all three occupation phases and compared with modern air temperature from Borgo Valsugana (TN; Extended Figure 3).

### Amelogenin analysis: sex estimation

Ibex sex was assessed on 21 adult specimens through the identification of amelogenins (AMELX and AMELY) in tooth enamel. The method was set up and tested on 3 modern *Capra ibex* with known sex (two males and one female) from the Gran Paradiso National Park, part of the Bones Lab archaeozoological collection. Sample preparation and protein extraction were performed at the Bones Lab of the Department of Cultural Heritage in Ravenna (UNIBO) and at the Department of Chemical and Geological Sciences (UNIMORE). For each individual, about 100 mg of enamel was sampled using a dentist drill. After rinsing with MilliQ water in an ultrasonic bath, the enamel chunks were leached for 5 minutes in 200 μL of 5% HCl and washed again with MilliQ. Samples were left overnight at room temperature in 750 μL of 1.2M HCl to extract enamel proteins. The solution was then collected and refrigerated. Enamel chunks were soaked again in 750 μL of 1.2M HCl and left overnight at room temperature. The two HCl solutions were then combined and the peptides were extracted and purified using C_18_ in house stage tips following the protocol for faunal samples described in^74^. Samples were finally measured by a LC-MS/MS, housed at the Centro Interdipartimentale Grandi Strumenti of the University of Modena and Reggio Emilia, using a Dionex Ultimate 3000 UHPLC coupled to a high-resolution Q Exactive mass spectrometer (Thermo Scientific, Bremen, Germany) (as in^75^). A Top5 DDA mode was selected with no inclusion list. Raw data were searched against a fasta reference dataset generated ad-hoc including dental proteome sequences from UniProt, using both Mascot and MaxQuant. Enzymatic digestion was set to ‘unspecific’ and the following variable modifications were included: oxidation (M), deamidation (NQ), and phosphorylation (ST). The final sex estimation, based on bovine reference sequences (entries AMELX_BOVIN P02817 and AMELY_BOVIN Q99004), is a combination of the results from the two searches. The detailed proteomic data analysis is described in Supplementary Note 4, which includes the estimates of deamidation and peptide lengths.

### Ancient DNA workflow

A total of four teeth for each phase of the Riparo Dalmeri rockshelter was selected for the genetics analysis, along with additional samples from Riparo Cogola (n = 2) and Romagnano Loc III (n = 2) to provide a broader baseline for the genetic variability of prehistoric ibex in the Alpine range. The ancient DNA workflow was conducted in the ancient facilities of the Department of Cultural Heritage (University of Bologna). The aDNA laboratory is physically separated from other laboratories of the institute and is pressurised to reduce air-influx, and equipped with UV-light to minimise exogenous DNA contamination. During the analyses, strict ancient DNA authenticity criteria were followed to support the authenticity of the results^76,77^. Sampling for the analyses was performed on the teeth: the surface of each sample was smoothly cleaned with 4% HCl, rinsed in 80% EtOH and then sterilised under UV-light for 20’. Before extraction, approximately 100 mg of dentine’s powder was collected in an Eppendorf tube 2 ml drilling the bone, with a precision drill (Dremel 8200). To reduce friction, we used a dental bit at a low rotation rate. The samples were extracted following the procedure using a silica-based protocol^78^ modified as in^79^. Molecular concentration was initially measured on a Qubit fluorometer and DNA extracts were used for single-stranded DNA libraries construction following the protocol of^80^. The libraries were purified with the MinElute PCR purification kit (Qiagen, Hilden, Germany) and quantified using Agilent 2100 Bioanalyzer DNA 1000 chips. Eleven out of twelve libraries reported good molecular content and were pooled in equimolar amounts for shotgun sequencing on HiSeqX Ten 2×75bp lane.

### Bioinformatic analysis for genetic data

We processed raw genetic data using EAGER v.2.5.1^81^. Forward (AGATCGGAAGAGCACACGTCTGAACTCCAGTCAC) and reverse adapters (AGATCGGAAGAGCGTCGTGTAGGGAAAGAGTGTA) were clipped, allowing filtering for clipped read length <30, minimum read quality ∼20 and minimum adapter overlap ∼1. We evaluated DNA preservation by mapping reads to the *Capra hircus* genome assembly (GCA_001704415.2_ARS1.2) using the bwa *aln* algorithm, allowing for a minimum number of mismatch ∼0.01, maximum edit distance ∼2 and length of seeds ∼1024. We filtered out reads with mapping quality <30 and removed duplicates with *markduplicates*. To reduce bias in genotyping from non-UDG-treated reads, we clipped off 5 bases from left and right ends of single-stranded mapped sequences. Post mortem damage patterns were calculated using DamageProfiler, with length filter ∼100 and number of bases for each read to be considered ∼30. Summary statistics were automatically generated by EAGER using multiqc, qualimap and samtools. We reconstructed two mitogenomic datasets by using EAGER v.2.5.1 again to process and map newly generated and available data from the literature against the *C. hircus* mitochondrial reference genome (NC_005044.2) and the *Capra ibex* mitochondrial reference genome (NC_020623). For the ancient samples, we used the mapDamage *rescale* option to downscale base quality for those positions of the mapped files that were affected by deamination patterns. Modern samples were directly reconstructed from full length, deduplicated .bam files, along with the samples produced in 2020 by Robin et al., which underwent UDG treatment. We used Mutect2^82^ to call variants from the bam files, and bcftools^83^ to normalise and filter the resulting variants by applying a minimum allele frequency (AF) > 0.5 for all samples, and minimum allele depth (AD) coverage > 3 for low coverage samples (n=12) and AD > 5 for all other samples. Consensus mitogenomes for each individual were reconstructed using bcftools consensus command.

#### Mitochondrial DNA investigation

##### Phylogenetic Analysis

To investigate the phylogenetic relationships of *Capra ibex* from Riparo Dalmeri, we generated three distinct mitogenome datasets: “Capra_all”, “Capra_ibex/pyrenaica”, and “Capra_ibex” (Supplementary Table S3). These datasets were constructed by combining new sequences from the six specimens obtained in this study with previously published mitogenomes of various Capra species, along with five sheep individuals used as an outgroup^6,7,84,85^. Only samples with mean coverage ≥ 10x were retained to build the datasets. Due to the difficulty of unambiguously identifying small remains of Alpine ibex compared to other *Capra* species based on morphology, we first constructed a maximum likelihood tree using the dataset *Capra_all*, to confirm the species of our samples. The software IQTree v2.0.3 (https://doi.org/10.1093/molbev/msaa015) was used for the tree with the *-m TESTNEW* command to search the best-fit model (TIM2+F+I+G4 chosen according to BIC). We performed a nonparametric bootstrap analysis with 1000 replicates and a search for the best-scoring maximum likelihood tree. Results were graphically visualised with FigTree^86^ and edited in Adobe Illustrator. To validate the decision to include only samples with a coverage ≥ 10x, the same tree (Supplementary Figure 3) was constructed using the mitogenome dataset “Capra_all_low”, with all available samples (Supplementary Table S3). Quality assessment of the low coverage mitogenomes from Cogola, Romagnano and Riparo Dalmeri is provided in the Supplementary Text (Supplementary Note 3).

To conduct a more in-depth analysis of the relationships among the *C. ibex* samples and to estimate the divergence times between Alpine ibex and Iberian ibex, we performed Bayesian inference using the “Capra_ibex/pyrenaica” dataset. We ran the analysis with the software BEAST v2.7.1^87^ and the BEAST package bdsky v1.4.5^88^. Default parameters were used except for the root node, which was set to 20000 years as the starting point for the birth-death skyline serial prior (the root node must be older than the oldest sample, which is 13400 cal. BP in this case). The package BEAST Model Test^89^ was used to find the best substitution model, and mutation rate estimation was performed under a strict clock assumption. A Markov Chain Monte Carlo (MCMC) run with 2,5×10^7^ generations was conducted, sampling every 1,000 steps. The chain convergence and effective sampling size (ESS) values were evaluated using Tracer v1.7.2^90^. The first 10% trees were discarded as a burn-in and the Maximum Clade Credibility tree was obtained using TreeAnnotator v2.7.1. Results were graphically visualised with FigTree^86^ and edited in Adobe Illustrator.

Haplotype diversity among the Alpine ibex from Riparo Dalmeri was explored building a TCS network from the dataset “Capra_ibex”. The network was constructed using PopArt^91^. We used the incorporated TCS algorithm, that calculates an absolute pairwise distance matrix of all haplotypes and connects them on the base of the parsimony criterion to minimise mutation steps between haplotypes. The resulting network was refined in Adobe Illustrator.

##### Genetic distance and mitochondrial diversity

To better explain the Network results, using the dataset “Capra_ibex”, we calculated the average pairwise distances for all ancient, historic and modern samples of *Capra ibex*, assuming they belong to distinct lineages. Also, we categorised the samples from the same dataset into five groups based on mitochondrial haplogroups and computed the within-groups mean distances for them to compare the genetic diversity for the different populations within *C. ibex*. Distance calculations were conducted using MEGA X^92^, with gaps and missing data handled as pairwise deletions to preserve the maximum number of homologous sequences. To analyse the mitochondrial genetic diversity, we used the dataset “Capra_ibex”. First, we divided the sequences in four age-sample groups: Dalmeri (13500 cal. BP to 11500 cal. BP), ancient (9700 cal. BP to 3500 cal. BP), historic (950 cal. BP to 1919 CE), and modern individuals. Subsequently, for each group, we calculated the number of haplotypes (Nh), haplotype diversity (h), segregation sites (S) and nucleotide diversity (π) with the R v4.4.0 software packages Pegas v1.3^93^.

##### Demographic analyses

To infer the changes in population history of Alpine ibex over the last 30000 years, we used a Bayesian Skyline coalescent prior applied to the mitogenomes^94^. To obtain the skyline plot we used the dataset “Capra_ ibex” and performed the analysis with Beast v.2.7.1^87^. The package BEAST Model Test^89^ was used to estimate the best substitution model, with a mutation rate of 2.73e^−7 95,96^ under a stick clock. We ran a MCMC chain of 2.5×10^7^ samples and visualised the MCMC chain convergence using Tracer v1.7.2^90^.

### Genetic sex determination

We used EAGER with the same parameters listed above to map shotgun sequences to the reference genome sequence of the Saanen domestic goat (GCA_015443085.1), the only available *Capra hircus* genome at the time of writing which presented annotated sex chromosomes. We tested sex assignment using the so-called “Mittnik approach”^97^, modified according to^98^ (see here: https://www.ncbi.nlm.nih.gov/pmc/articles/PMC7144076/). We combined the “Skoglund approach”^99^ with the approach presented in^100^ to validate our results. Detailed explanation can be found in Supplementary Note 3.

### Data availability

Raw sequencing data are publicly available in the European Nucleotide Archive under project number PRJEB87623 (data is private pending evaluation). All the proteomic raw data, the MaxQuant evidence file, and the deamidation R script were uploaded to Zenodo (https://doi.org/10.5281/zenodo.15024003). Isotope data are available within the manuscript and its supplementary material. The OxCal code used for modelling the radiocarbon dates of Riparo Dalmeri was uploaded to Zenodo (https://doi.org/10.5281/zenodo.15063853).

## Acknowledgments

We thank the Soprintendenza per i Beni Culturali of the Autonomous Province of Trento, and the MUSE - Science museum of Trento for granting access to the archaeological materials analysed in this study. Dr. Diego Pinetti and Dr. Filippo Genovese of the Centro Interdipartimentale Grandi Strumenti (UNIMORE) are acknowledged for technical assistance during LC-MS/MS analyses. We also thank Prof. Luca Pagani (University of Padua) for his feedback and comments, which have provided helpful insights for this work. Nerobutto sponsored research activities on prehistory carried out at Muse - Science museum of Trento between 2019 and 2025. FL is supported by the European Union’s Horizon Europe Research and Innovation program under the Marie Skłodowska-Curie Actions PF (grant agreement no. 101104566—AROUSE). MR is supported by CHANGES, SPOKE 5 “Science and Technologies for Sustainable Diagnostics of Cultural Heritage,” PE 0000020, CUP B53C22003890006, NRP M4C2, Investment 1.3, funded by the European, Union— NextGenerationEU. SS is supported by the ERC FIRSTSTEPS project (Grant Agreement ID: 101019659).

## Extended Data

**Extended Figure 1.**
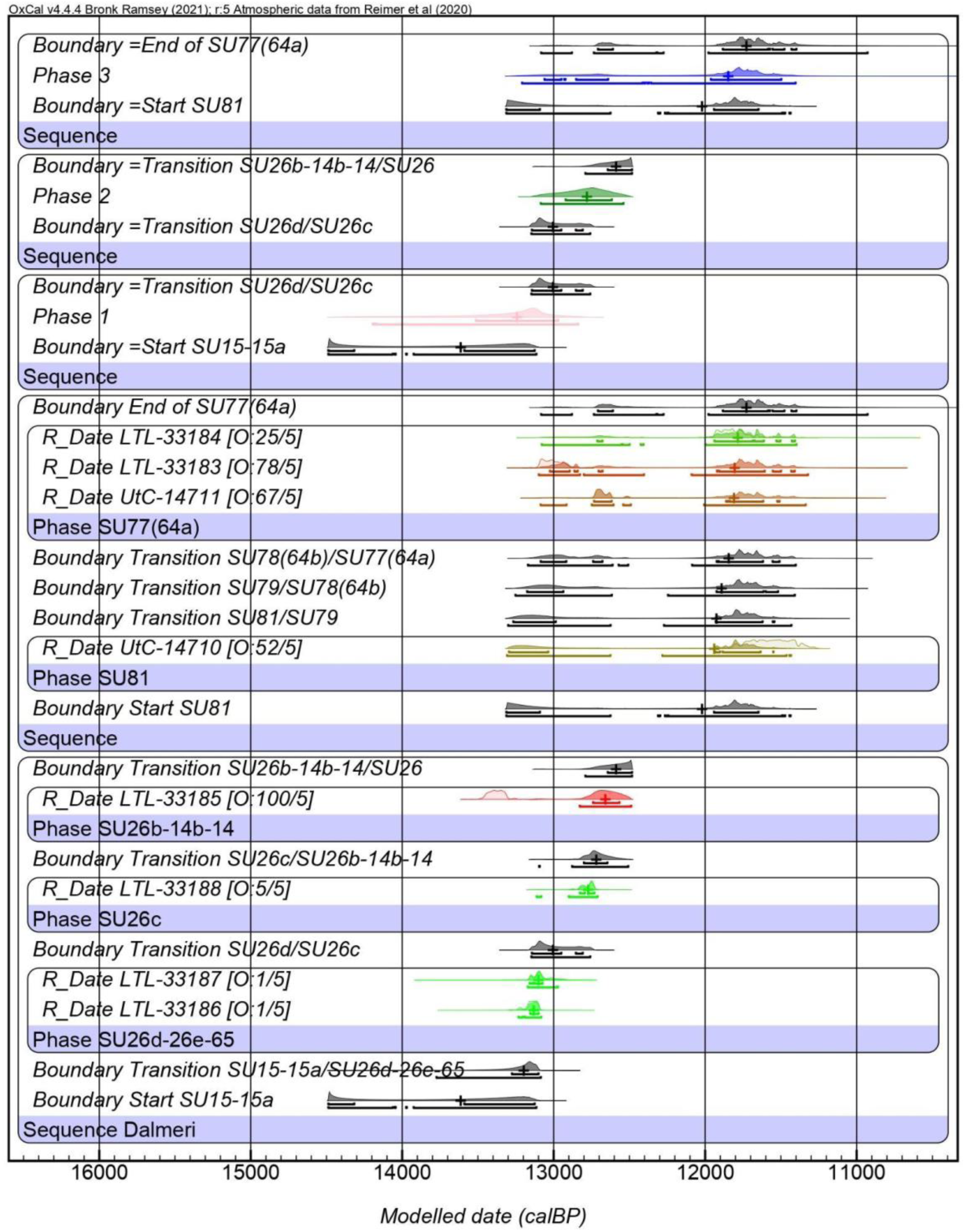
Bayesian modelled ^14^C dates from Riparo Dalmeri inner (1-2) and outer (3) frequentation phases calculated after direct radiocarbon analysis of bones and teeth samples.

**Extended Figure 2.**
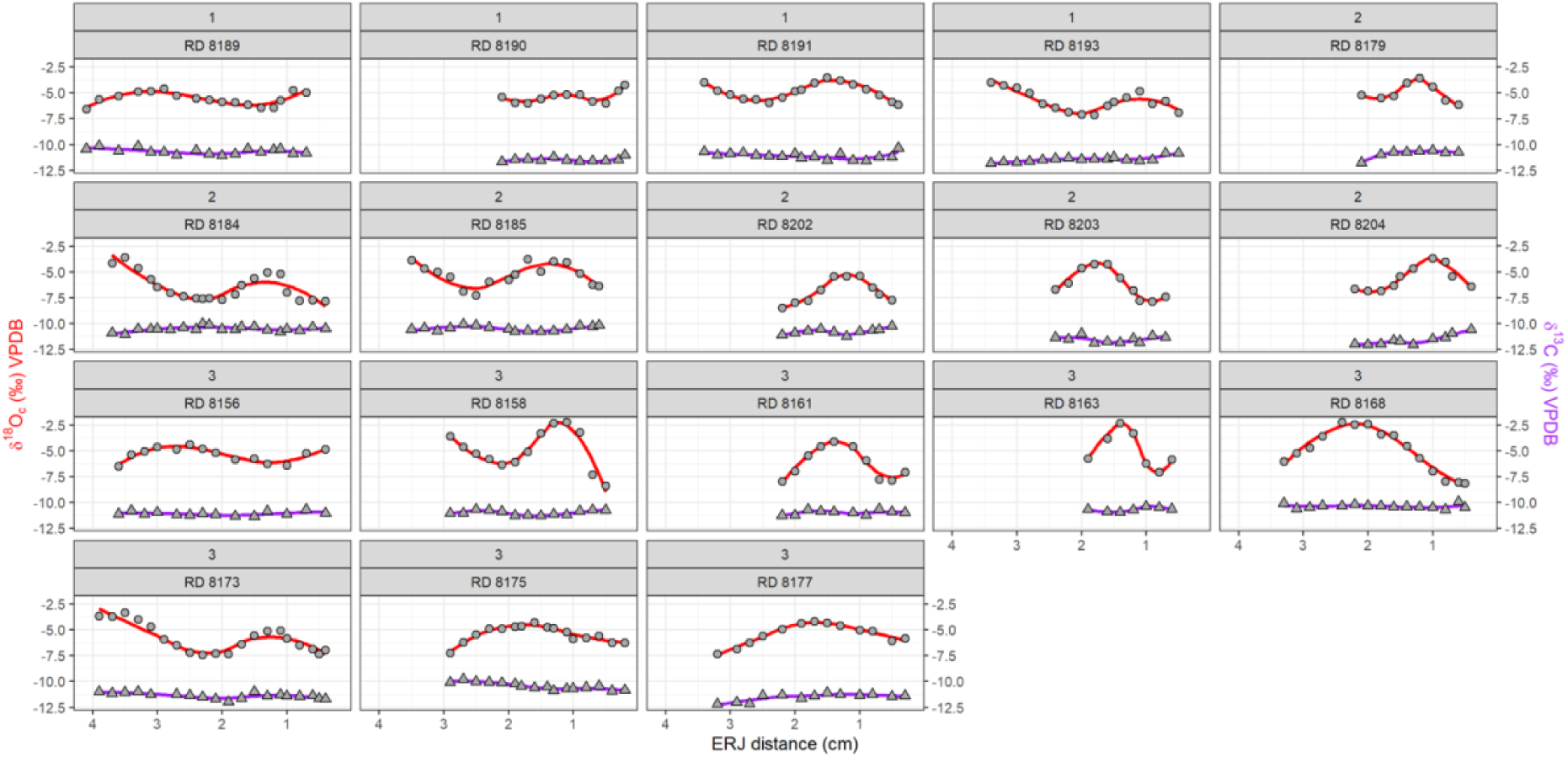
Intra-tooth δ^18^O and δ^13^C (‰ VPDB) profiles. Carbon isotope values (triangles, purple profiles) are homogeneous between samples, while oxygen isotope values (circles, red profiles) show sinusoidal patterns indicative of local climate fluctuations.

**Extended Figure 3.**
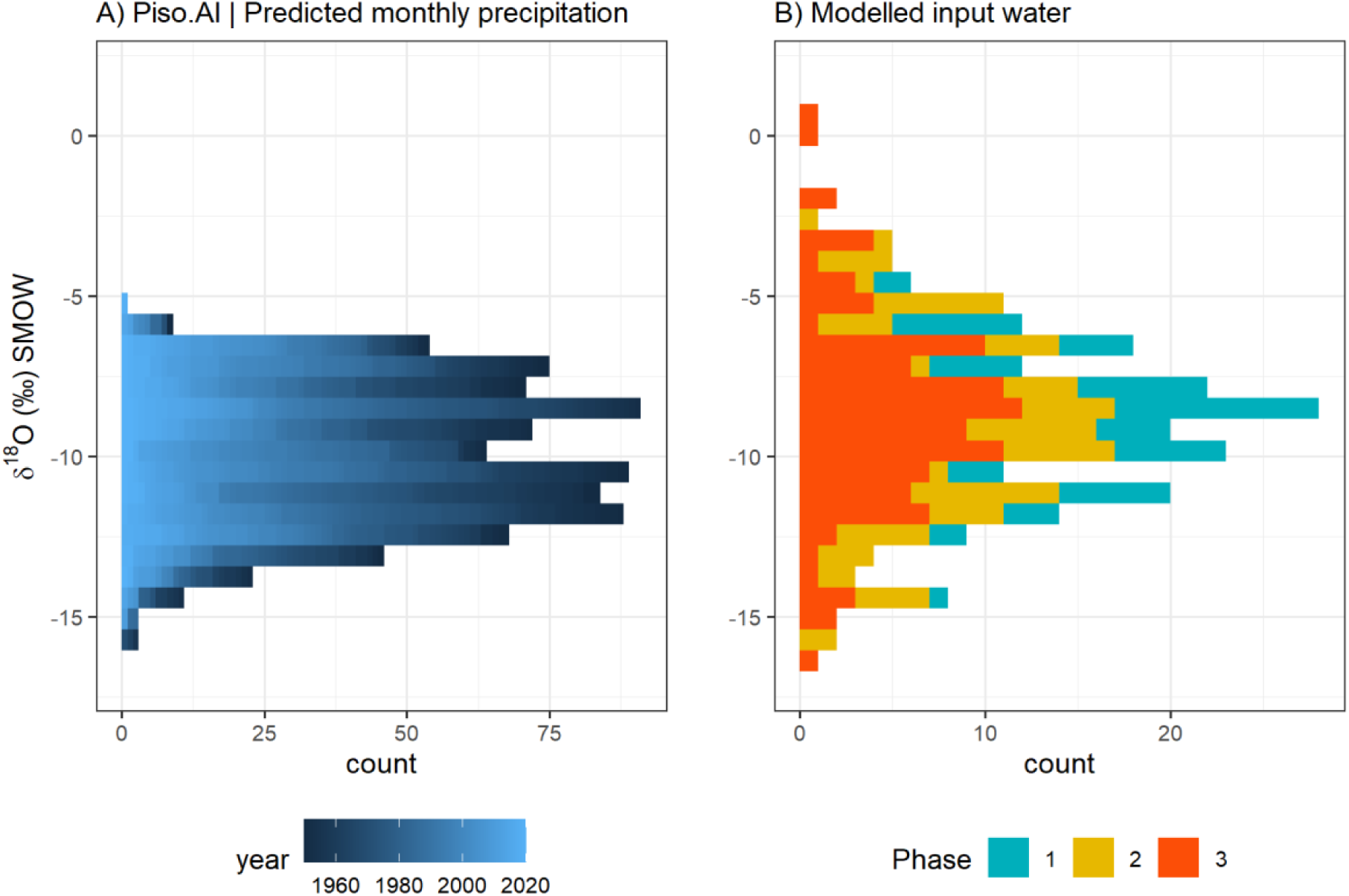
Comparison of **A)** modern predicted monthly time-series precipitation δ^18^O at the coordinates of Riparo Dalmeri (Piso.AI; https://isotope.bot.unibas.ch/PisoAI/) with **B)** modelled data δ^18^O (input water) from ibex teeth. In A) data from the last 70 years were reported.

**Extended Figure 4.**
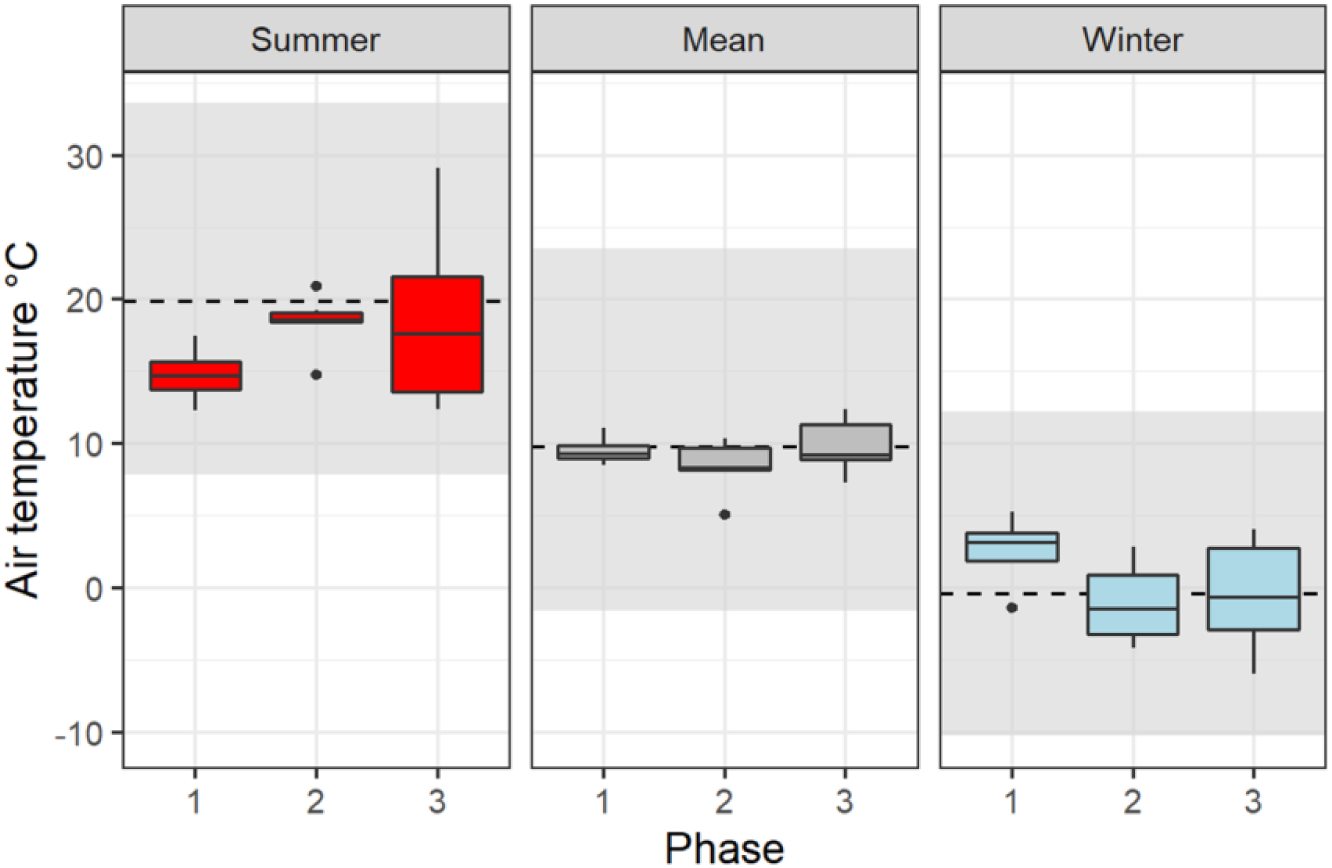
Summer, mean and winter air temperature estimation divided by phase, obtained from modelled input water oxygen isotope data of ibex enamel. The dashed lines represent mean for summer, winter and mean annual temperatures in Borgo Valsugana (TN). Gray areas are the maximum-minimum ranges for summer, winter and mean annual temperatures in Borgo Valsugana.

**Extended Figure 5.**
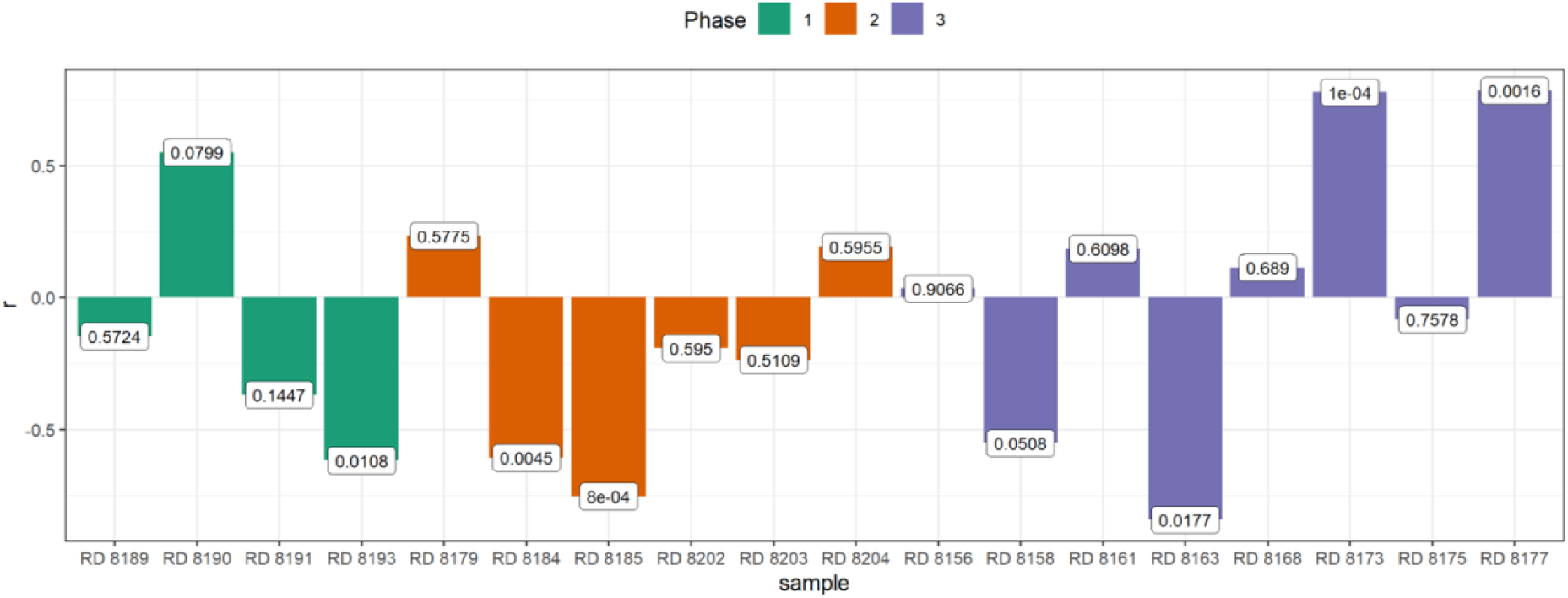
Linear correlation between δ^18^O and δ^13^C values of ibex teeth (n = 18) divided by site occupation phase. The correlation coefficient r > 0.5 indicates a strong positive correlation (i.e., C-O covariation), while r < 0.5 indicates a strong negative correlation (i.e., C-O anti-covariation). Labels are p-values.

**Extended Figure 6.**
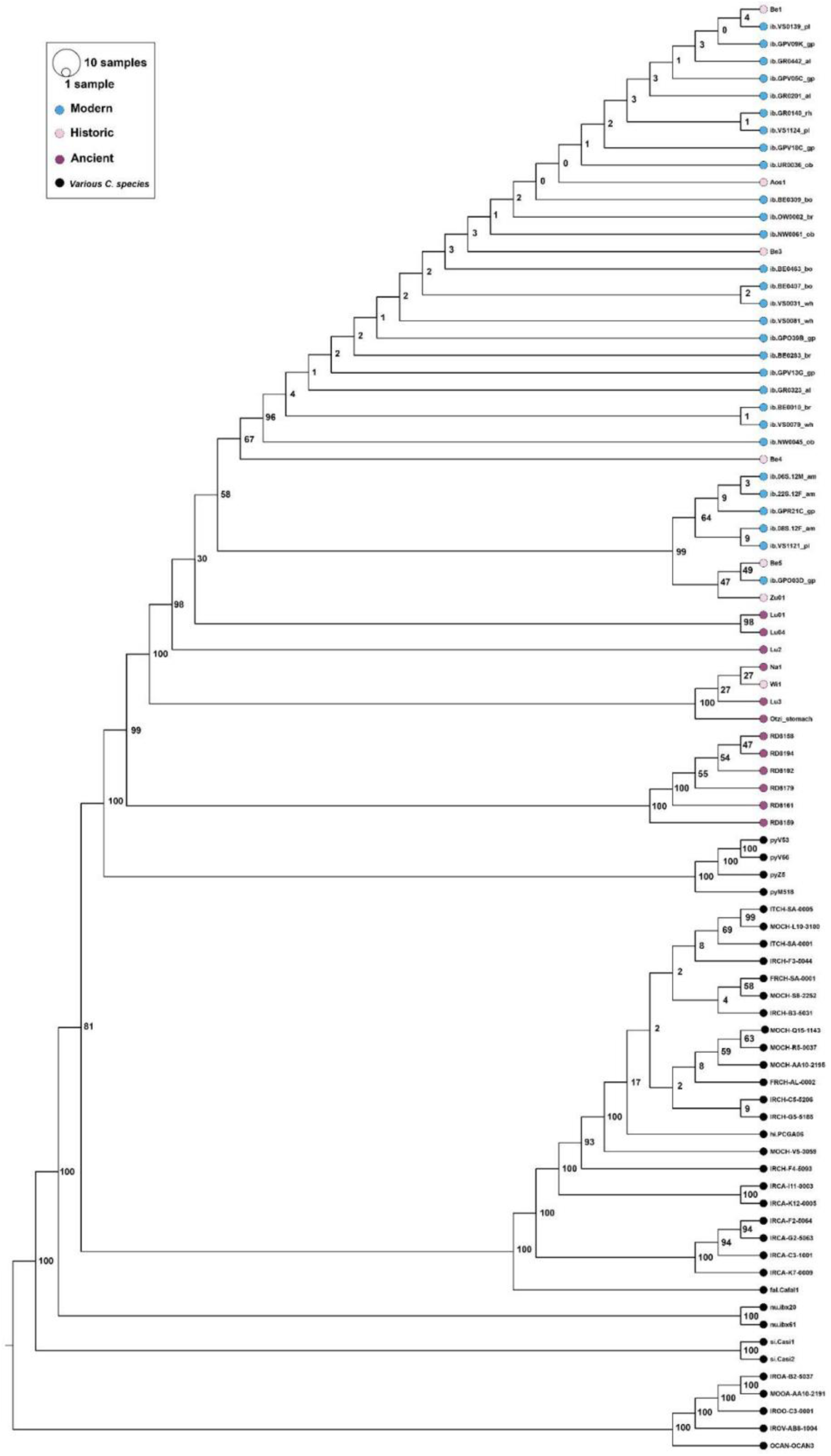
Maximum likelihood tree with high-coverage samples.

**Extended Figure 7.**
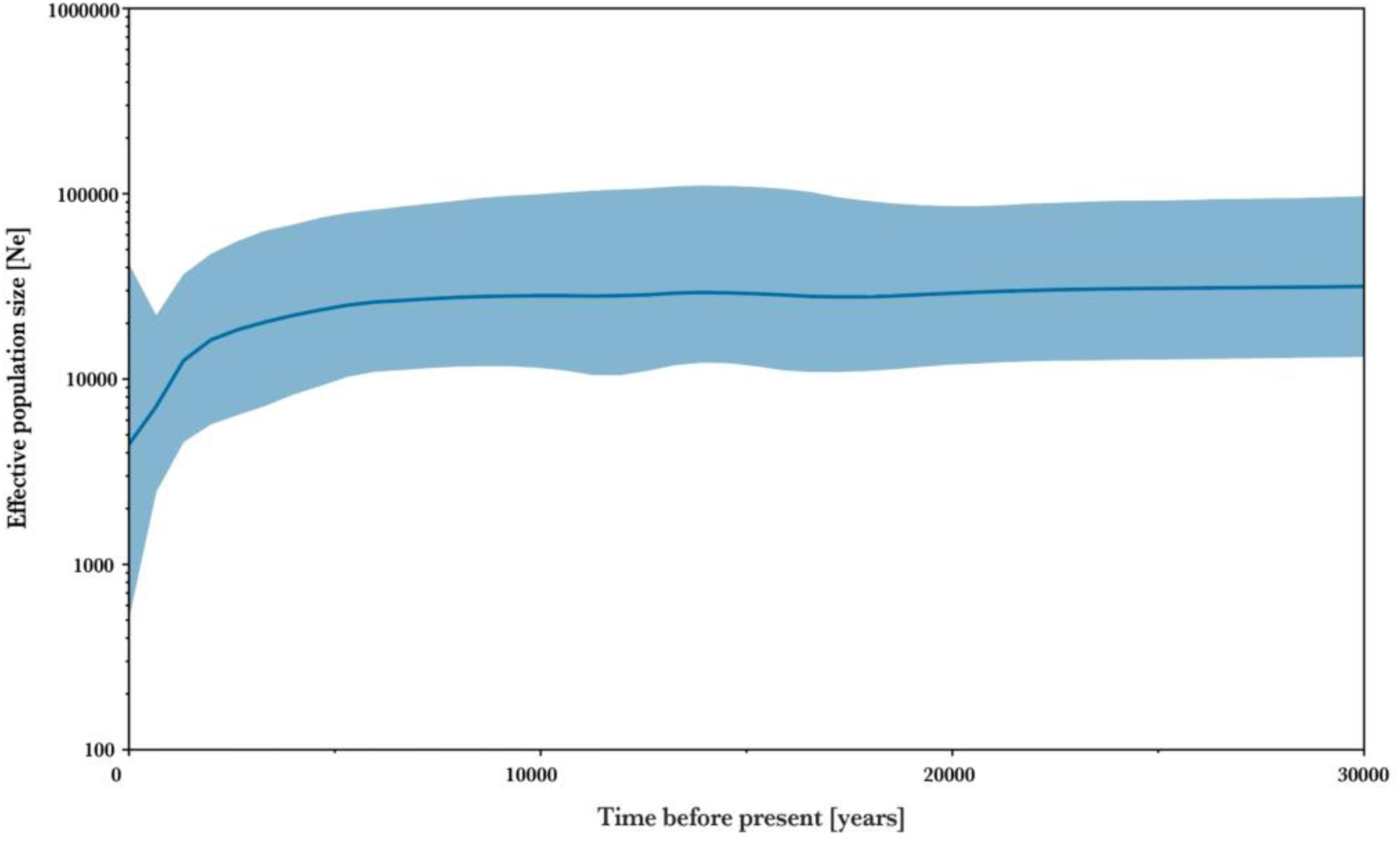
Bayesian skyline plot based on the *Capra ibex* dataset. A mutation rate of 2.73e−7 under a strict clock model was used, with an MCMC chain of 25 million samples. The x-axis represents time, while the y-axis represents the population size of *Capra ibex* expressed as Ne.

**Extended Figure 8.**
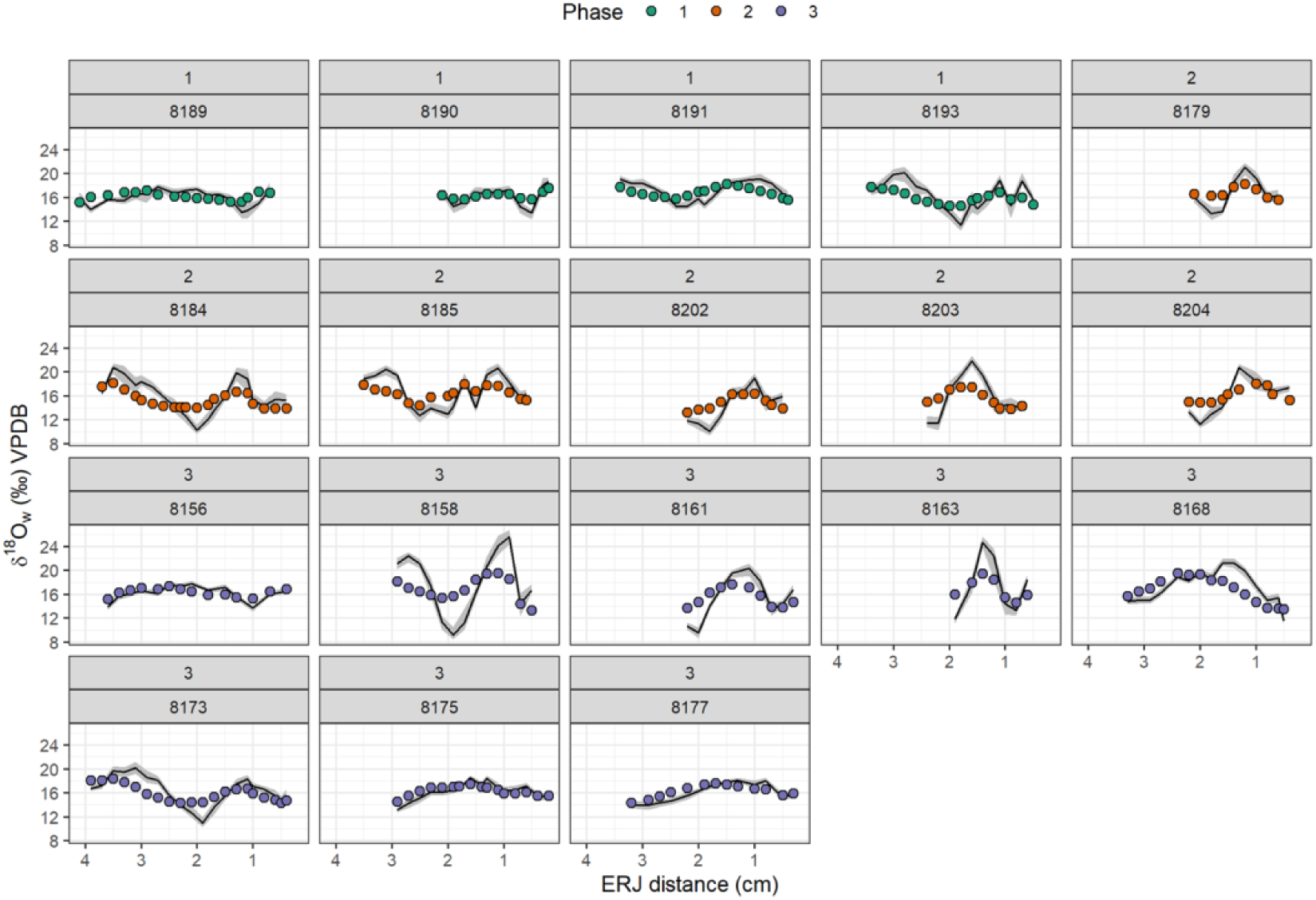
Modelled input δ^18^O_water_ (‰ VPDB) profiles. Circles are the Dalmeri δ^18^O values converted into ingested water values^73^ and coloured according to the occupation phase. Black lines are the environmental input data obtained by inverse modelling^67^. The grey ribbons represent the maximum and minimum estimations from the model. The distance measured from the enamel-root junction (cm) is on the x-axis.

**Extended Table 1.** Zooarchaeological material from Riparo Dalmeri.

| *Sample ID* | *US* | *Phase* | *Species* | *Tooth class* | *Age class* |
| --- | --- | --- | --- | --- | --- |
| RD_8168 | 21 | 3 | C. ibex | M3 | Young adult |
| RD_8185 | 26b roof | 2 | C. ibex | M3 | Adult I |
| RD_8173 | 26 cleaning | 3 | C. ibex | M3 | Young adult |
| RD_8191 | 26d | 1 | C. ibex | M3 | Adult I |
| RD_8202 | 26c | 2 | C. ibex | M3 | Adult II |
| RD_8203 | 26c roof | 2 | C. ibex | M3 | Adult II |
| RD_8204 | 26c roof | 2 | C. ibex | M3 | Adult II |
| RD_8193 | 26d | 1 | C. ibex | M3 | Adult I |
| RD_8158 | 64 | 3 | C. ibex | M3 | Adult I |
| RD_8161 | 64a | 3 | C. ibex | M3 | Adult II |
| RD_8190 | 26d | 1 | C. ibex | M3 | Adult II |
| RD_8179 | 26b | 2 | C. ibex | M3 | Adult II |
| RD_8163 | 64b | 3 | C. ibex | M3 | Adult I |
| RD_8189 | 26d | 1 | C. ibex | M3 | Adult I |
| RD_8156 | 77 | 3 | C. ibex | M3 | Young adult |
| RD_8184 | 26b roof | 2 | C. ibex | M3 | Adult I |
| RD_8175 | 26 | 3 | C. ibex | M3 | Young adult |
| RD_8177 | 26 | 3 | C. ibex | M3 | Young adult |
| RD_8159 | 64a | 3 | C. ibex | M3 | Senile |
| RD_8192 | 26d | 1 | C. ibex | M3 | Adult II |
| RD_8194 | 26d | 1 | C. ibex | M3 | Adult II |
| RD_8205 | 14b | 2 | C. ibex | p2 | Young adult |
| RD_8206 | 14b | 2 | C. ibex | p3 | Young adult |
| RD_8210a | 14b | 2 | C. ibex | p3 | Young |
| RD_8210b | 14b | 2 | C. ibex | p4 | Young |
| RD_8210c | 14b | 2 | C. ibex | M1 | Young |
| RD_8210d | 14b | 2 | C. ibex | M2 gem | Young |
| RD_8212 | 26 | 3 | C. ibex | i4 | Young |
| RD_8214 | 14b | 2 | C. ibex | p4 | Very young |
| RD_8215 | 14 | 2 | C. ibex | p4 | Young |
| RD_8218 | 26c | 2 | C. ibex | p4 | Young |
| RD_8202 | 26c | 2 | C. ibex | M3 | Adult II |
| RD_8230 | 64b | na | Lepus timidus | Incisor | na |
| RD_8220a | 26 | na | Micromammal | Incisor | na |
| RD_8219 | 26 | na | Micromammal | Incisor | na |
| RD_8224 | 26 | na | Micromammal | Incisor | na |

**Extended Table 2.**
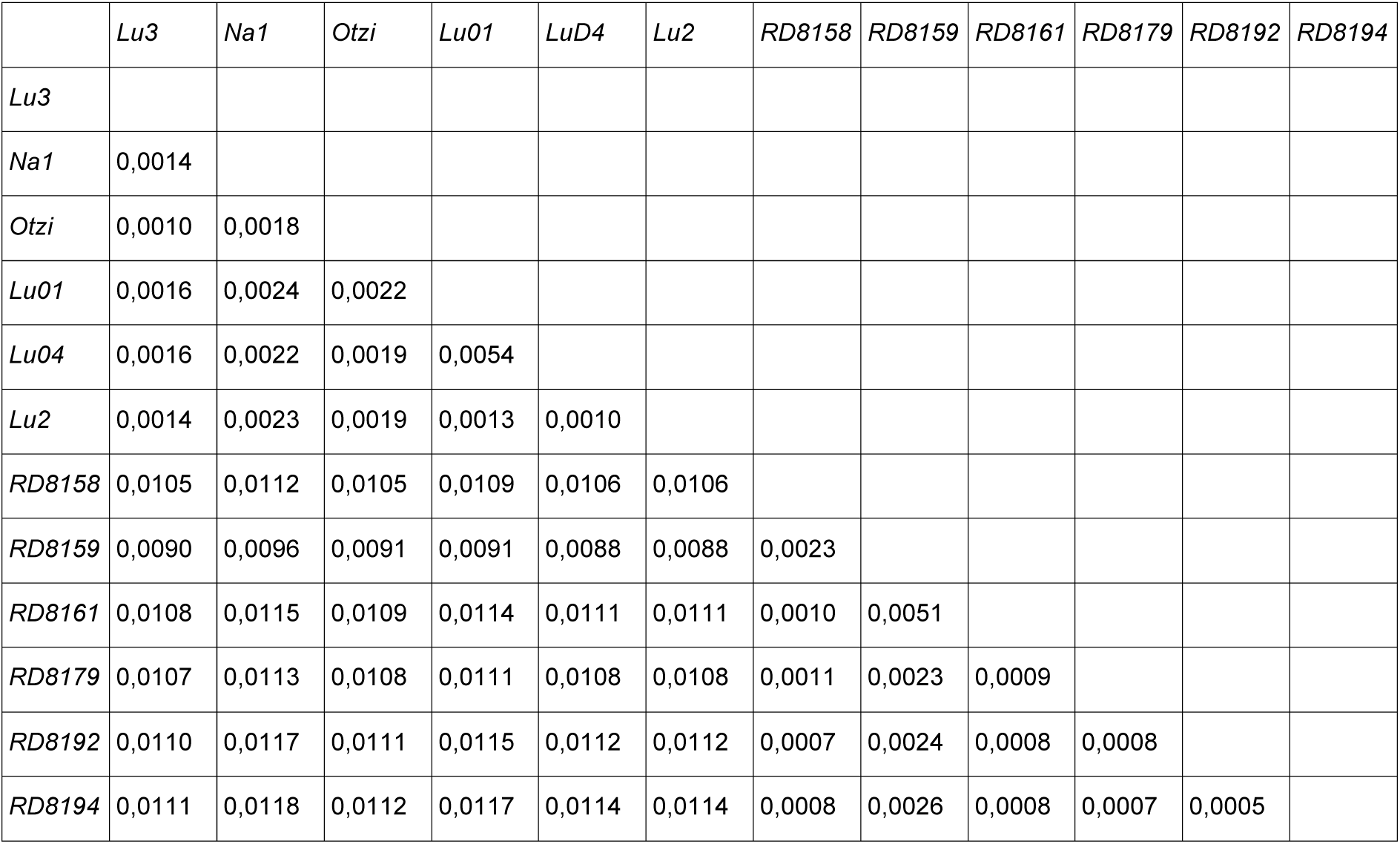
Pairwise distance for all ancient Capra ibex samples.

