## Supplementary Information for "Ecology and demographic structure of an extinct ibex population in Upper Palaeolithic Italian Alps"

### Supplementary Text

#### Supplementary Note 1: Archaeological information

*(Duches, R., Fontana, A., Nannini, N., Romandini, M., Terlato, G.)*

##### Late Pleistocene human adaptations in the Alps

The peopling of northeastern Italy after the Last Glacial Maximum was a gradual process marked by the progressive colonization of new territories that had been previously abandoned, following changes of vegetation and animal distribution[1,2]. Late Epigravettian penetration, firstly limited to valley floors and high plateaus around 500 m a.s.l.[3,4], reaches mid-altitude territories during the second part of the Late Glacial interstadial with the full development of a logistical occupation network at the limit between coniferous woods and alpine prairies[5]. This organization reflects a seasonal mobility strategy that sometimes involved sites with complementary functions, located at different altitudes. Specifically, Late Epigravettian groups established semipermanent occupations on valley floors, characterized by the functional division of settlement spaces (e.g., workshops for flintknapping, dwelling structures, butchering areas, etc.) and mid-altitude seasonal camps where specialized tasks, such as hunting ungulates from the Alpine prairie (mostly *Capra ibex*), meat processing, and hide and flintworking were carried out. During the Younger Dryas, climatic and environmental changes had a significant impact on Epigravettian societies[6–8], leading to several transformations in the subsistence strategies of these groups. The size and complexity of the sites suggest, in fact, the existence of higher mobility patterns compared to the previous period[9,10]. Technological analyses suggest that lithic production systems gradually simplified their structure throughout the Late Glacial interstadial, indicating a shift in technical investment from core shaping to shaping of the derived flake blank[11–13]. On the other hand, a persistence of standardized lithic backed tools used as projectile implements is attested throughout the Alleröd[11,14]. This persistence is thus a result of flexibility in retouching, framed in a progressively simplified production system. Such adaptive technology must have been encouraged by Late Glacial climatic and environmental changes and the occupation of previously inaccessible alpine territories. Thus, it can be argued that the flexibility of technical behaviors represents a key factor in the transformation of Late Epigravettian societies throughout the Late Glacial, enabling them to adapt and evolve in response to environmental, social and economic changes[11].

#### Riparo Dalmeri

Since its discovery in 1990, many studies have outlined a complex and exceptional picture of this rock shelter, one of the most important Palaeolithic sites in Northern Italy[15]. Among the seasonal sites in north-eastern Italy, Riparo Dalmeri is the only one that can be considered truly specialized in ibex hunting[16]. In addition to the greater abundance of *C. ibex* remains (see Supplementary Note 2) its uniqueness lies in its characteristics of a base camp (e.g., dwelling structures, hearths, tools, ornaments and portable art), where all the family members – including children[17] – lived seasonally basing much of their subsistence on ibex[18,19]. Paleoenvironmental reconstruction indicates an open alpine prairie with some emerging wooded areas of pines and larches nearby[5,20]. The presence of species other than ibex in the faunal record, including other ungulates, small mammals, carnivores, birds and fishes indicates that the territory exploited by hunter-gatherers was quite large, extending from alpine meadows to the valley floor of the Brenta River[21,22]. However, ibex was the most hunted species and the main reason for site occupation. The great importance this animal had to Epigravettian hunter-gatherers is also stressed by its representation on some painted stones and the presence of pits intentionally filled with ibex cranial bones and horns. These findings seem to be related to a complex ritual that could broaden the interpretation of the site to the symbolic and ritual sphere, at least in the first phase of occupation[15,19,23].

The site, located at about 1240 m a.s.l., overlooks the head of a small periglacial valley, a tributary of the deep Valsugana canyon (Trentino) crossed by the Brenta River. The morphogenesis of the rockshelter results from the differential erosion of the stratified limestone bedrock (oolitic lithofacies of the Jurassic Rotzo Formation), caused by the combined effects of carbonate dissolution and cryogenic processes during the Last Glacial Maximum[20,23]. The shelter faces N-E and extends NNW-SSE for 30 m. The rock overhang extends up to 7 m, with an actual height of 4 m above the ground. The shelter's stratigraphic sequence spans from the end of the Upper Pleistocene to the Holocene. It is approximately 4.5 m thick and, in the inner part of the shelter, it has been divided from bottom to top into the following stratigraphic complexes[15,23,24]:

1) <u>The stratigraphy preceding the settlement phase</u> consists of a sequence of thermoclastic breccias, indicative of generally cold and humid conditions. Specifically, an early deposit associated with carbonate silts of karst and/or colluvial origin was identified directly on the top of the bedrock. Upward the breccia is enriched with a matrix of aeolic origin. Finally, an open-work breccia with clasts characterized by thin capping, typical of climatic conditions involving frost. SU 50 is distinguished by its enrichment in a clay matrix of karst and/or colluvial origin.

2) <u>The first phase of human occupation</u> of the shelter involves the creation of a structure near the current line of the overhang. It is represented by an accumulation of collapse

blocks (SU 74), at the top of which was found a large stone with an anthropomorphic figure (RD 211). This structure partially covers a breccia with an organic matrix (SU 65), which has a strong anthropogenic component (e.g., lithic industry, faunal remains, and charcoal) associated with numerous ochre-painted stones (n = 267). Unit 65 constitutes a lens that thins toward the interior of the shelter, while reaching a maximum thickness of 45 cm near the present overhang. Most of the painted stones are found within this unit, while some are scattered in the innermost part of the shelter, resting on the roof of a cryoclastic breccia (SU 15a). Overall, the distribution of stones forms a band of about 30 $m^2$, more than 4 meters wide, with an east-west orientation, oblique to the alignment of the inner rock wall. Human occupation during this phase is further supported by the presence of two hearths in the east part of the shelter and the remains of a dwelling structure with a 4 m diameter.

3) The later Epigravettian dwelling floors (SU 26c, 26b/14b) are characterized by the superposition of Ah-type organic horizons related to anthropogenic activity. These horizons developed on a parent material consisting of a cryoclastic breccia (US 15a), enriched by silty, micaceous, blackish, highly organic frank matrix with abundant anthropogenic remains. The two main archaeological horizons 14-26b and 26c, in stratigraphic succession, were excavated over an area of 84 $m^2$ and preserved a substantial amount of lithic industry, industry on bone, organic animal remains, human decidual teeth, ornamental objects, and engraved stone tool cortex. The identified structures allowed the settlement area to be subdivided into a western section, where the sub-circular hut was still in use, and an eastern section, where hearths were located. Within the hut, an accumulation of faunal remains and an area showing traces of burning were identified.

4) The upper sequence of the shelter consists of stratified breccias divided into 4 subunits, primarily supported by a silty matrix, with some areas exhibiting complete or partial clastic support. The matrix consists of biogenic “moonmilk”, whose deposition is linked to the onset of temperate climatic conditions, characterized by increased precipitation and temperature, typical of the Holocene Climatic Optimum. In the inner part of the shelter, the attribution of these layers to the Holocene is not supported by either radiocarbon dating or tecno-typological analysis of the lithic industry. Although the latter has only been studied preliminarily, the assemblage appears consistent with a Late Epigravettian industry, with some distinct features related to the end of this chrono-cultural complex[10,14,25]. Specifically, the presence of

geometric microlithic armatures and trapezoidal bitruncations suggests that these layers date to the Younger Dryas or the Pleistocene-Holocene transition.

The external succession of Riparo Dalmeri is divided into several units. Some of these units are heterotopic to those found in the inner part of the shelter, while others are of more recent age[26]. Above the occupation layers dating to the Late Glacial interstadial, the external succession shows a stratigraphic gap that roughly corresponds to the Younger Dryas. Sedimentary accumulation resumed at the Pleistocene-Holocene transition, culminating in a younger phase of sedimentary input around the Preboreal-Boreal shift[26]. More specifically, the lower part of the external succession consists of coarse clastic sediments with significant anthropogenic inputs (units 83, 84, 85, 85a, 88, 89, and 92, structure 86), which represent the lateral, heterotopic continuation of the habitation surfaces found in the inner portion of the stratigraphy. Above these units, the succession includes units 64, 77 (formerly 64a), 78 (formerly 64b), 7,9, and 81. At the outermost position, the sequence consists of units 85, 82, 76, 75, 9,1 and 90, listed from bottom to top.
The complex of units 64, 77, 78, 79, and 80 forms a sigmoidal lenticular deposit with lateral interfaces (inward and outward from the shelter) inclined at approximately 45°, with the upper interface sub-horizontal and the lower interface inclined at 10-15° outward from the shelter. The complex of units 90, 91, 75, and 76 forms a sequence of layers with biconcave geometry, while the basal complex, composed of units 82, 83, and 85, consists of tabular layers slightly inclined toward the exterior of the shelter[26].
Regarding the younger phase of sedimentary input in the external succession (units 64, 77, 78, 79, and 80), inconsistencies have emerged between radiocarbon dating and chrono-cultural attribution based on lithic industry. Previous dates obtained from charcoal[26] suggested a younger phase of human occupation around the Preboreal-Boreal shift. However, the lithic assemblage is not consistent with an attribution to the First or Early Mesolithic[27,28]. Since no evidence was found to confirm the existence of Mesolithic human frequentations, it is possible that these layers resulted from the reworking of pre-existing anthropogenic material from the Late Epigravettian. Therefore, previous Holocene dates obtained on charcoal may reflect the time of paleosol formation rather than the primary deposition of anthropogenic material.

#### Riparo Cogola

The rockshelter site of Riparo Cogola is located 1070 m a.s.l. on the northern edge of the Sette Comuni Plateau, on the Vicentine Alps (Trento, NE Italy). Excavations conducted between 1999 and 2002 by researchers from the Museo Tridentino di Scienze Naturali identified three main periods of frequentation: the most ancient during the Final Epigravettian (SU 19, radiocarbon dated between 12600-11950; 12900-12450 cal. BP); a transitional phase between the Epigravettian and the

Sauveterrian (SU 18, radiocarbon dated at 11250-11150 cal. BP); and the Sauveterrian (SU 17-16, the latter radiocarbon dated 10750-10550 cal. BP)[20]. In all three phases, the primary hunting prey consisted of ungulates (red deer, roe deer, ibex, and chamois), while wild boar was present only in the first two phases and elk only in the Early Sauveterrian. The analysis of the skeletal remains indicates that ibex carcasses were taken into the shelter whole and then butchered, as evidenced by striae on the lithic tools and impact marks. In contrast, a selective butchering process was applied to deer carcasses, with only the anterior and posterior portions being brought inside the shelter. Studies on the hunting-season of ibex and chamois show they were killed in summer-autumn period, suggesting that humans occupied the shelter during the warmer months[29]. The two samples selected for paleogenetic analysis were directly radiocarbon dated following the same procedure described in the Materials and Methods section. Results align with the anthropic frequentation phases of the rockshelter, with sample 0830 radiocarbon date 12650-12500 cal. BP ($1\sigma$) and sample 0831 radiocarbon dated 11150-10800 cal. BP ($1\sigma$).

#### Romagnano Loc III

The rockshelter site of Romagnano Loc III was discovered in 1968 during quarrying activities. Located at 210 m a.s.l., approximately 10 km south of Trento (NE Italy), the site hosts an extremely rich and well-preserved archaeological deposit, spanning from the Mesolithic to the Iron Age[28,30–34]. The Mesolithic deposits were excavated between 1971 and 1973 by researchers from the Museo Tridentino di Scienze Naturali and the Università di Ferrara[35]. Several layers were identified and systematically radiocarbon dated from 11703 to 7280 cal. BP[28,36]. The faunal assemblage collected from the Middle and Recent Sauveterrian levels AC primarily consists of ungulates, with a significant prevalence of *Capra ibex* and *Cervus elaphus*. The two *C. ibex* samples analyzed in this paper (ID 6123 and 6124) were found in layer AC8. This stratigraphic unit was dated to 10500-10250 cal. BP and was associated with Middle-Sauveterrian lithic technology.

### Supplementary Note 2: Zooarchaeological data

*(Duches, R., Fontana, A., Nannini, N., Romandini, M., Terlato, G.)*

Zooarchaeological studies conducted on the faunistic assemblage from Riparo Dalmeri[16,19,21,37–42] have provided valuable insights into the site's economy and clarified the methods of animal exploitation. The analysis involved more than 120,000 bones, with over 15,000 determined to species or genus level (Supplementary Table S5). In all phases, hunting was primarily focused on ibex (*Capra ibex*), with a lesser emphasis on red deer (*Cervus elaphus*). Wild boar (*Sus scrofa*), roe deer (*Capreolus capreolus*), elk (*Alces alces*), and chamois (*Rupicapra rupicapra*) are represented by very few remains. Among carnivores, the most numerous remains belong to the bear (*Ursus arctos*), followed by fewer elements attributed to the fox (*Vulpes vulpes*), wolf (*Canis lupus*), and badger (*Meles Meles*). Remains of the hare (*Lepus sp.*), beaver (*Castor fiber*), marmot (*Marmota marmota*), and European hedgehog (*Erinaceus europaeus*) are exceptionally rare. Fish remains were also identified, with Cyprinids comprising 90% of the total. The majority of these remains belong to the barbel (*Barbus plebejus*) and the chub (*Leuciscus cephalus*), while fewer are attributed to trout (*Salmo Trutta*) and grayling (*Thymallus thymallus*). Pike (*Esox lucius*) remains are extremely rare. The presence of individuals measuring around 30/40 cm supports the hypothesis that fishing was the primary method of capture[43]. The bird remains (n = 146) are mainly attributable to Galliformes, including Tetraoninae, such as the black grouse (*Tetrao tetrix*), Lagopedes (rock ptarmigan, *Lagopus cfr. Mutus*, and willow ptarmigan, *Lagopus cfr. Lagopus*) and small Phasianidae (common quail, *Coturnix coturnix*). Passeriformes are also relatively abundant. Most of these birds were undoubtedly introduced into the shelter by humans, as indicated by anthropogenic traces, particularly on the Galliformes remains[37].

Hunting of ungulates was primarily focused on early-stage adults and older adults, likely to maximize meat yield. The seasonality of human frequentation, determined by the presence of ungulate teeth from very young and young individuals and confirmed through the analysis of tooth thin sections, indicates that hunting occurred between summer and autumn[19,39,40,42,44]. Butchering marks are abundant both on ibex and red deer bones. Zooarchaeological data, based on the comparative frequencies of recovered skeletal remains, reveal a differential transport of animal carcasses from the initial kill site to the rockshelter. Hunters typically brought entire carcasses of medium-sized prey to the shelter, while only specific portions of larger animals, such as red deer, were selected for transport. Consistently, the frequency of ibex skeletal portions indicates that all elements of the carcass are represented, with a balanced distribution between the right and the left halves. The scarcity of some elements is attributed to differential bone preservation and the high rate of fragmentation, which in turn results from intense anthropogenic exploitation of the carcass. The spatial distribution of animal remains in the 26c anthropogenic horizon revealed a concentration of bones exhibiting cut marks and percussion marks. This area was therefore dedicated to the processing and butchering of ibex carcasses. The fracturing of bones took place inside the shelter, as evidenced by the higher frequency of percussion cones compared to the impact points found on the diaphyses. The presence of many percussion cones and vertebral discs suggests that the shelter was systematically cleared of larger fragments[40]. Additionally, the concentration of burnt remains points to areas where fires were frequently lit, with bones also used as fuel. Rare gnawing marks and a few bones attributed to bear and wolf cubs indicate that carnivores alternated with hunters in occupying the shelter.

Results for length, width, and thickness measured on third molars (M3) Riparo Dalmeri are summarized in Supplementary Table 1. The results are compared with data published in[19] (Supplementary Table 2).

**Supplementary Table 1**. Max anterior-posterior diameter (MAP) and transversal diameter (TD) measurements (in mm) for M3 teeth in the Dalmeri specimens. Age classes: JAd (young adults), Ad I (early-stage adults), and Ad II (older adults).

| *ID* | *TD* | *MAP* | *sex* | *Age* |
|---|---|---|---|---|
| RD_8153 | 10,3 | 27,8 | NA | Ad II |
| RD_8155 | 10 | 27,2 | NA | Ad II |
| RD_8157 | 9,8 | 27 | NA | JAd |
| RD_8158 | 9,7 | 28,2 | Female | Ad I |
| RD_8161 | 9 | 26,1 | Female | Ad II |
| RD_8164 | 8,48 | 27,4 | NA | JAd |
| RD_8165 | 9,32 | 29,8 | NA | Ad I |
| RD_8166 | 9,09 | 28,8 | NA | Ad I |
| RD_8170 | 9,6 | 29,2 | NA | JAd |
| RD_8171 | 9,6 | 27 | NA | Ad I |
| RD_8172 | 8,9 | 24,4 | NA | Ad I |
| RD_8175 | 10,2 | 28,1 | Female | JAd |
| RD_8179 | 10 | 28 | Male | Ad II |
| RD_8180 | 10,1 | 28,3 | NA | Ad I |
| RD_8181 | 10,1 | 29,2 | NA | Ad I |
| RD_8182 | 10,2 | 27,9 | NA | JAd |
| RD_8183 | 10,3 | 30,9 | NA | Ad I |
| RD_8184 | 10,2 | 27,7 | Male | Ad I |
| RD_8185 | 10 | 29,4 | Male | Ad I |
| RD_8187 | 10,6 | 29 | NA | Ad I |
| RD_8200 | 9 | 27,6 | NA | Ad II |

| | | | | |
|---|---|---|---|---|
| RD_8201 | 10,3 | 29,5 | NA | JAd |
| RD_8202 | 10,6 | 27,6 | Male | Ad II |
| RD_8203 | 10,1 | 26,2 | Male | Ad II |
| RD_8204 | 10 | 28,7 | Female | Ad II |

**Supplementary Table 2**. Comparison of MAP measures of lower M3 registered in several sites and reported in[19].

| *Site* | *n. samples* | *Max* | *Min* |
|---|---|---|---|
| Grotta del Broion* | 30 | 32,2 | 23,4 |
| Grotta di Fumane* | 7 | 30,7 | 28,1 |
| Paglicci Str.21-28* | 54 | 33,6 | 27,4 |
| Cala Str.Q* | 2 | 28,1 | 27,4 |
| Paglicci Str.6-1* | 6 | 30 | 24 |
| Tagliente* | 6 | 32,4 | 26 |
| Dalmeri (this study) | 25 | 30,9 | 24,4 |
| Dalmeri* | 14 | 30,59 | 26,29 |
| Romagnano III* | 6 | 27,3 | 23 |
| Cogne (modern populationi)* | 7 | 24,5 | 23 |

The graph in Supplementary Figure 1 shows that the median MAP value for Dalmeri is 28 mm, which falls within the measurement range observed at other sites, spanning from 23 mm to 33,6 mm. This range highlights a notable degree of variability in M3 dimensions. The comparison suggests differences in dental size between older and more recent populations. Specifically, the M3 dimensions from Dalmeri are comparable to those from the Broion site, but exhibit variations when compared to more recent populations, particularly those from Romagnano III and modern ibex specimens. The results suggest some variation in M3 dimensions based on both age and sex. However, no significant correlation is observed between the measurements, age, and sex of the individuals from Dalmeri. Adult males generally exhibit larger M3 dimensions than females, though some overlap is present (Supplementary Figure 2).

**Supplementary Figure 1.** MAP median values of lower M3 teeth from Dalmeri, compared with other sites spanning from the Middle Paleolithic to the early Holocene, as well as some modern ibex specimens. Data of other sites are from[19].

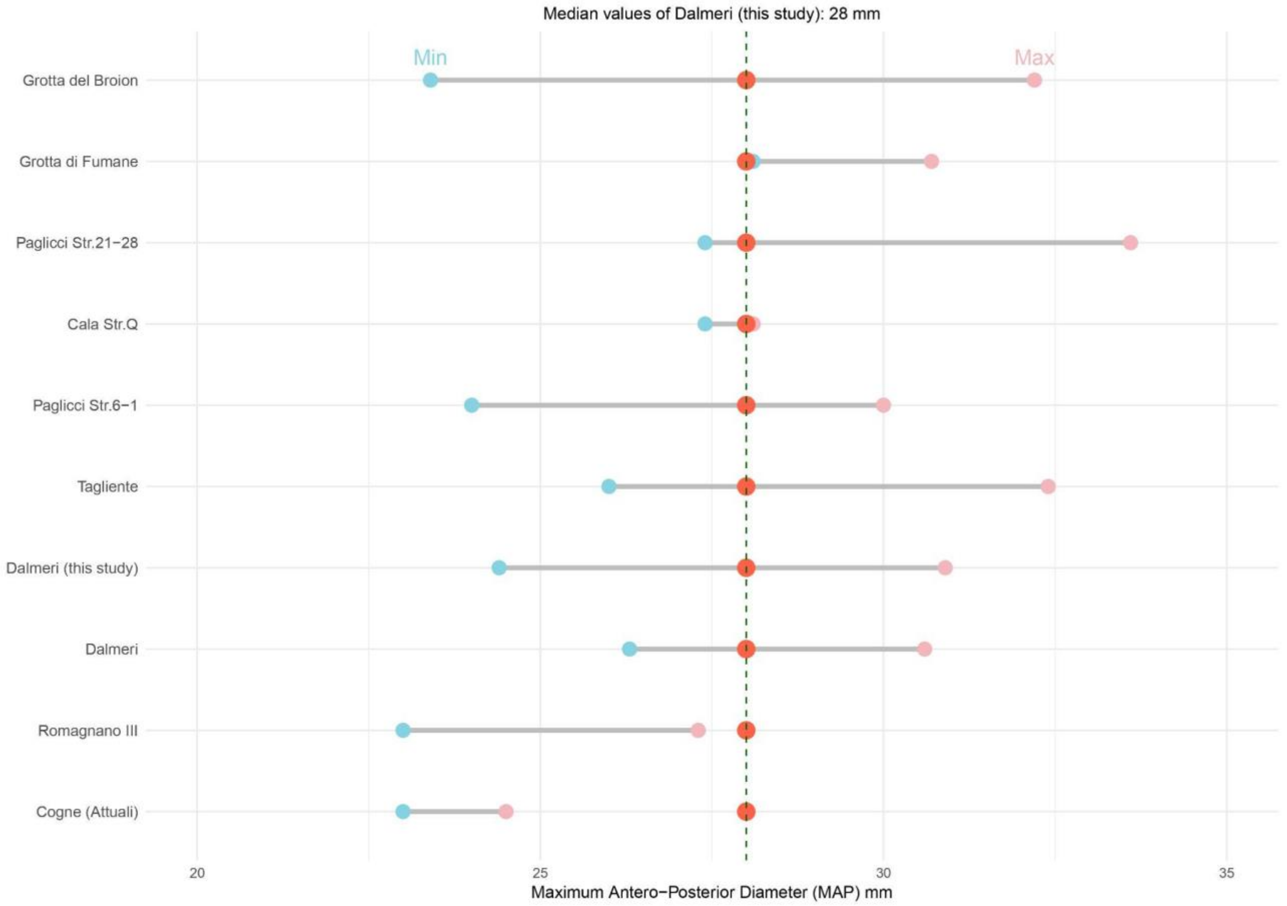

**Supplementary Figure 2**. Scatter plot of M3 measurements from Dalmeri, analyzing the relationship between age and sex of individuals using Kendall’s tau correlation. Maximum Antero-Posterior Diameter (MAP) and transversal diameter (TD) in millimeters. Age classes: JAd (young adults), Ad I (early-stage adults), and Ad II (older adults).

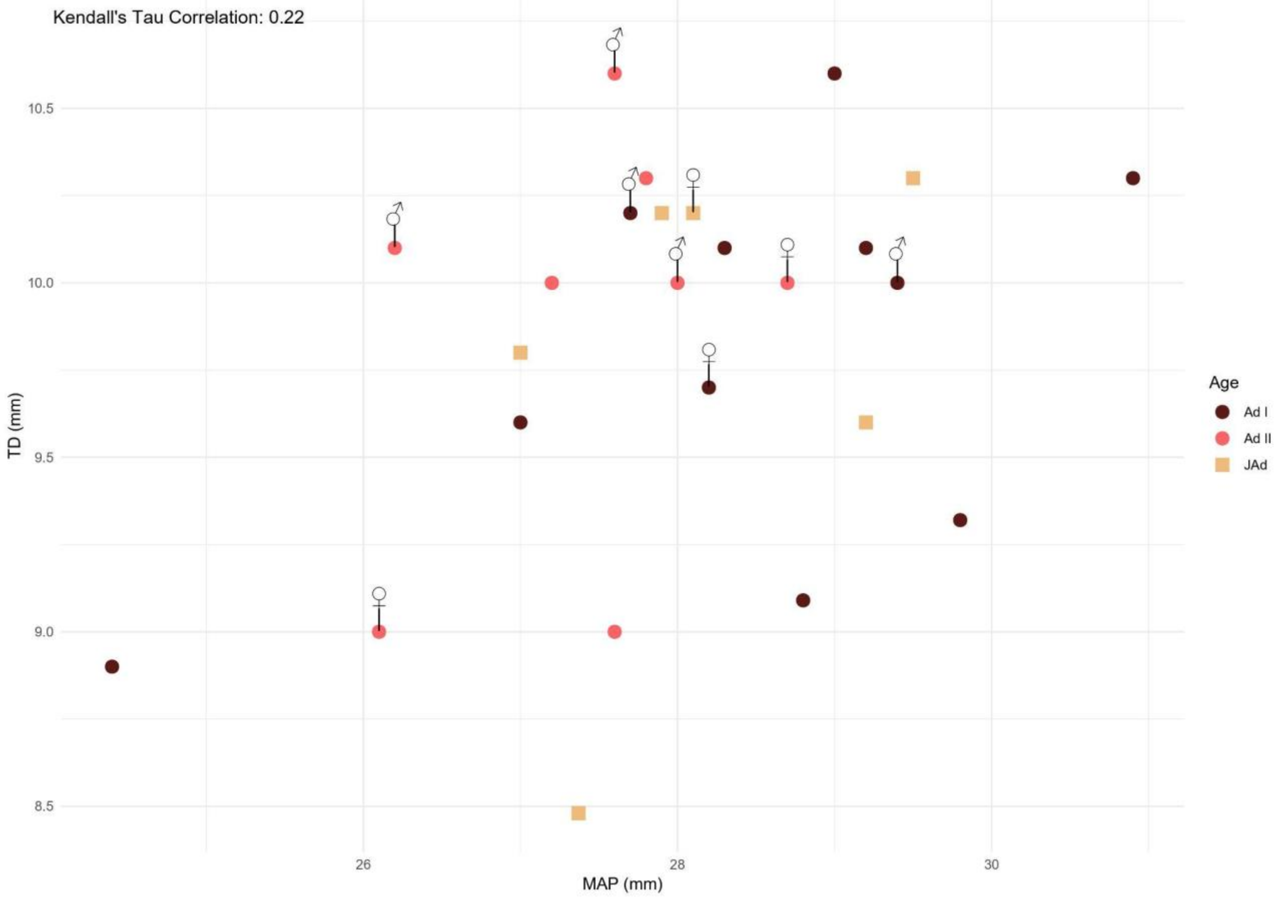

### Supplementary Note 3: Phylogenetic analysis of ancient mitogenomes

*(Fontani, F., Iacovera, R., Luiselli D., Cilli, E.)*

For phylogenetic analysis of high quality mitogenomes (>10x), we reconstructed three different datasets. All of the newly generated data from Riparo Dalmeri displayed distribution of C-to-T transition patterns in the first position at 5' ends ranging 30% to 47%, and the mean length of most mapped fragments spanned from 52 to 76 bp, indicating that the ancient DNA was authentic. The dataset "Capra_all" contained 48 *Capra ibex* samples (29 modern, 7 historic, 6 ancient, and 6 samples from Riparo Dalmeri), 4 *Capra pyrenaica*, 6 *Capra aegagrus*, 2 *Capra sibirica*, 1 *Capra falconer*i, 2 *Capra nubiana*, 16 *Capra hircus,* and 5 *Ovis aries*. For this dataset, we have removed the entire D-loop due to its high intraspecific variation in the control region, for a total of 15,529 bp. The dataset "Capra_ibex/pyrenaica" contained 44 *Capra ibex* (29 recent, 7 historic, 6 ancient and the 6 samples from Riparo Dalmeri) and 4 *Capra pyrenaica*. This dataset contained 15,603 sites of the 16,716 possible sites. We furthermore produced a third dataset "Capra ibex" composed of entire mitochondria, which only contained *Capra ibex* specimens. For each dataset, we used ClustalW/X[45] to align the sequences. All the historical and ancient mitochondrial sequences used to construct the three datasets have a mean coverage of ≥ 10x and yielded a robust phylogenetic tree Extended Figure 5). In fact, mitogenomes characterized by a coverage < 10x occupy unstable positions in the topology of the "low coverage" maximum likelihood tree (Supplementary Figure 3) computed from the dataset "Capra_all_low" (Supplementary Table S3). This instability not only results in questionable phylogenetic placements for these samples but also negatively impacts the statistical robustness of the associated nodes, as evidenced by significantly reduced bootstrap values. Including such low-coverage samples compromises the overall resolution of the tree, increasing uncertainty in the definition of phylogenetic relationships and invalidating the reliability of the analysis. Therefore, their exclusion is necessary to preserve the quality and consistency of the phylogenetic inference and subsequent analyses. Based on these criteria, five mitochondrial sequences from Riparo Dalmeri (RD_8163, RD_8190, RD_8202, RD_8203 and RD_8204), the two samples from Loc di Romagnano (6123, 6124), the two samples from Riparo Cogola (830, 831) and three mitochondrial sequences from Robin et al. 2022 (Gro1, Pil2 and Zue2) were discarded from the analysis.

**Supplementary Figure 3**. Maximum likelihood tree with low coverage samples.

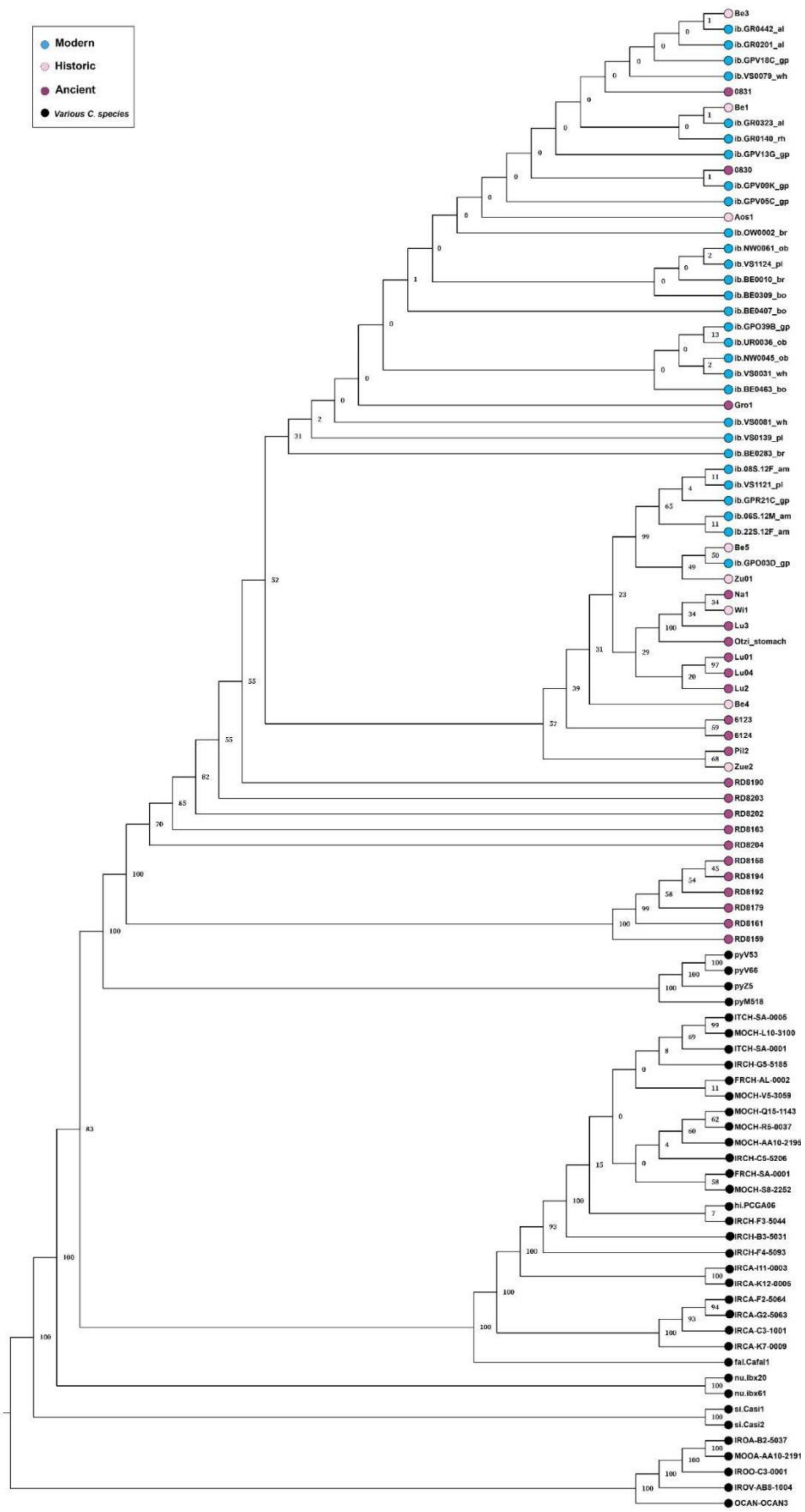

Interestingly, all the haplotypes reconstructed from the Dalmeri mitogenomes (haplogroup A) are unique, and no genetic affinity is visible between samples from the same archaeological phase. Haplogroups B and C, also made up only of unique haplotypes, consist of the remaining ancient samples, along with the outlier Wi1. The two remaining haplogroups consist exclusively of modern and historical samples, with the latter being the most numerous of the five (n = 27). Among them, nineteen modern individuals from both the south and north of the Alps belong to the most common haplotype. Conversely, only one modern Alpine ibex sample from the Swiss Eastern Alps shares its haplotype with three historical samples from Italy. All remaining historical samples in the haplogroups D and E carry unique haplotypes. As a consequence, average genetic distances for ancient ibex are about six times greater than that of historical ibex (0.6% compared with 0.1%), while those of modern ibex are just 0.04% indicating poor genetic diversity in current populations (Supplementary Table 3-4). The Alpine ibex specimen Ötzi_stomach shows a genetic distance ranging from 0.10% to 0.21% when compared to samples from the Lucerne region, which cover a time span from 9,679 to 3,457 cal. BP. A similar genetic distance of 0.17% is observed when Ötzi_stomach is compared to the Na1 sample (5,445 - 4,963 cal. BP), which is geographically close to Lucerne. Despite being territorially more similar to the Riparo Dalmeri samples, which cover a time span from 13,500 cal. BP to 11,500 cal. BP, Ötzi_stomach shows a mean genetic distance of 1.06% from them, which is identical to that observed between Riparo Dalmeri and the other ancient samples.

**Supplementary Table 3.** Average genetic distances. Measures among the different *Capra ibex* groups are based on 16,716 bp homologous mitochondrial sequences.

| *Group* | *Average genetic distance (%)* |
|---|---|
| Modern | 0,041 |
| Historic | 0,1 |
| All Ancient | 0,653 |
| Haplogroup A | 0,134 |
| Haplogroup B | 0,126 |
| Haplogroup C | 0,091 |
| Haplogroup D | 0,044 |
| Haplogroup E | 0,014 |

**Supplementary Table 4.** Haplotype and nucleotide diversity in ibex populations

| *Population* | *N. of samples* | *Haplotype number* | *Haplotype diversity* | *Segregation sites* | *Nucleotide diversity* |
|---|---|---|---|---|---|
| Modern | 29 | 14 | 0,869 | 31 | 0,0003 |
| Historic | 7 | 6 | 0,952 | 48 | 0,0009 |
| Ancient | 6 | 6 | 1 | 66 | 0,001 |
| Dalmeri | 6 | 6 | 1 | 57 | 0,001 |

### Supplementary Note 4: Paleoproteomics

*(Armaroli, E., Silvestrini, S., Lugli, F.)*

We collated a protein reference database containing entries for COL1A1, COL1A2, COL2A1, CO4A1, AMELX, AMELY, TUFT1, MMP20, KLK4, AMBN, ALB and ENAM of *Bos taurus*, *Sus scrofa*, *Ovis aries*, *Capra hircus*, *Capra ibex* and *Homo sapiens*, derived from UniProt. Proteomic data analysis was performed through MaxQuant (MQ) v2.1.0.0 using unspecific digestion and setting deamidation (NQ), phosphorylation (ST), and oxidation (M) as variable PTMs. Peptides' length was allowed to be between 7 and 25 amino acids, with a minimum score of 35, and filtered to FDR = 1% at peptide and protein level; other settings were left as default. Preliminary Mascot searches (vs. SwissProt) on modern *Capra ibex* specimens with known sex, indicated that UniProt reviewed amelogenin sequences of *Bos taurus* show the expected pattern among sexes (i.e., AMELY in male only; Supplementary Figure 4), while unreviewed and incomplete AMEL sequences of *Capra ibex*, *Ovis aries* and *Capra hircus* show worse coverage and no sex-related differences. We therefore decided to rely on AMEL entries for *Bos taurus* to estimate the sex of fossil samples. All individuals with at least one specific-AMELY peptide after MQ search were estimated as males, after validating this approach on the three modern samples (note e.g. that ibex_mod_M1 shows a single AMELY peptide and is a male); all the peptides were then pooled and their intensities summed (Supplementary Figure 5).

**Supplementary Figure 4**. Mascot fragmentation spectra of sexual-dimorphic AMELY (Q99004) and AMELX_BOVIN (P02817) peptides, obtained from modern *C. ibex* samples. No AMELY peptides were found in the modern female individual. The main sexual-dimorphic region observed is highlighted in yellow; sequences of AMELX_BOVIN and AMELY_BOVIN were aligned with Clustal O (1.2.4).

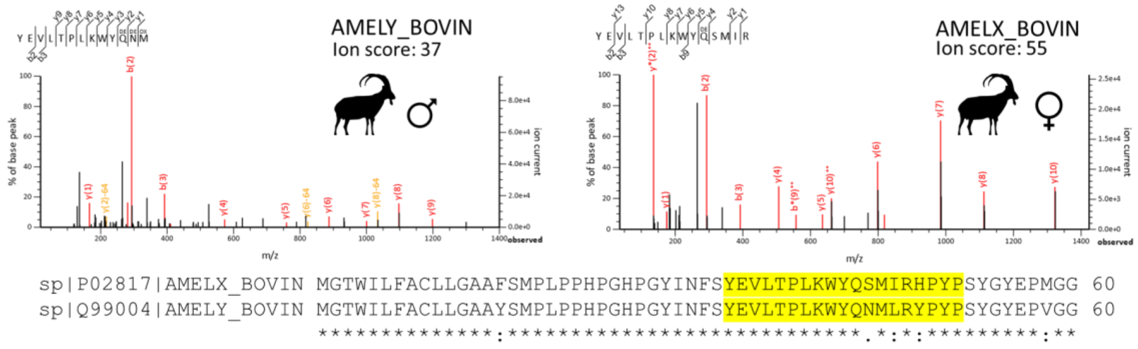

**Supplementary Figure 5**. Total intensities of razor-AMELX and AMELY peptides from fossil and modern (‘ibex_mod’) *C. ibex* enamel samples after MaxQuant search. Half-circles have an AMELY intensity equal to 0.

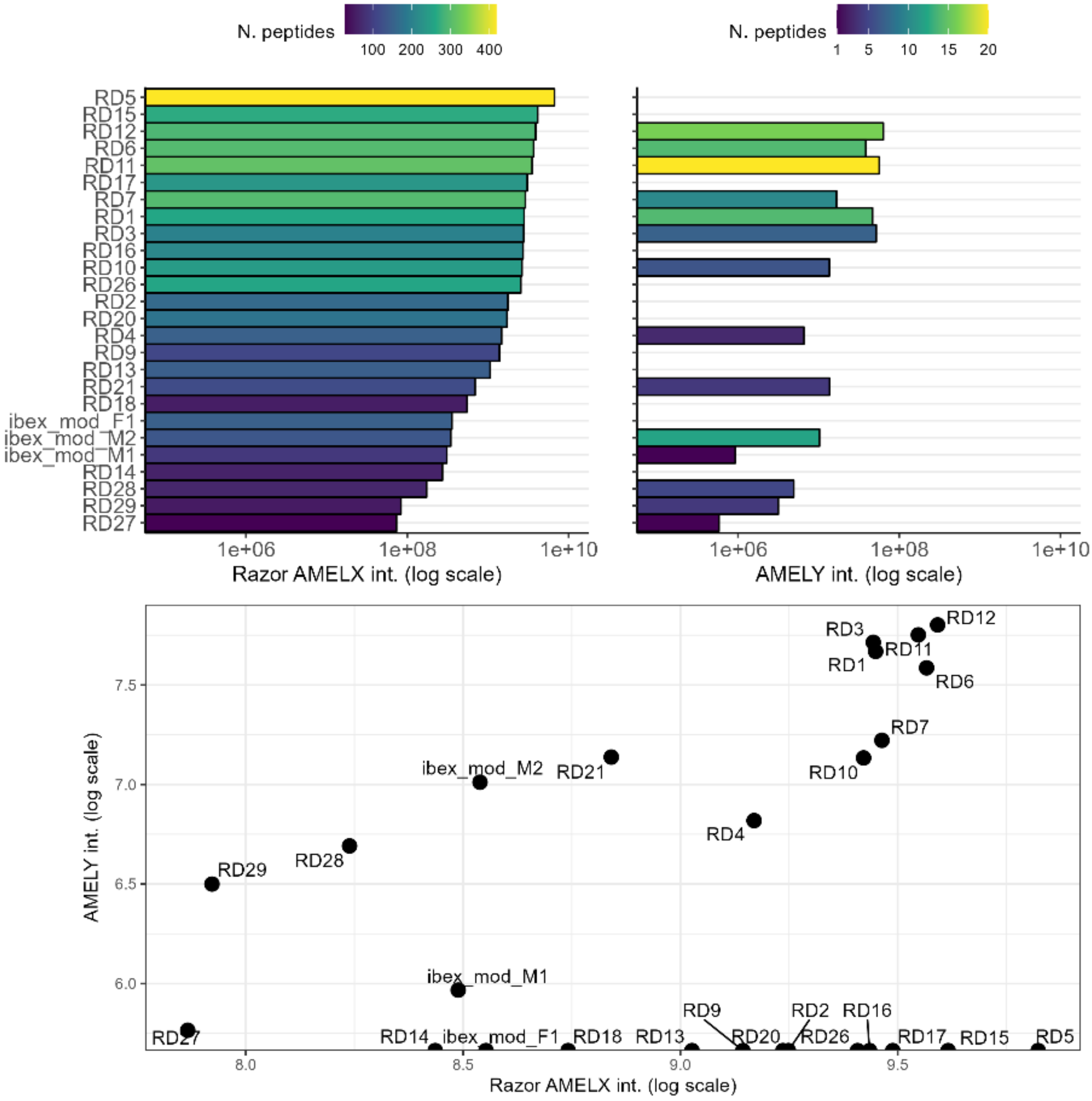


An initial comparison with DNA sexing indicated proteomic sex estimation of RD9 as female false positive, a known issue in amelogenin-based sex estimation; all the other individuals agreed with DNA. We thus decided to perform a second search run using Mascot for all those samples that resulted as females through MQ; the two search engines are indeed known to occasionally provide different matching peptides (e.g.[46]). For the Mascot search, we used the same PTMs, FDR threshold and reference fasta dataset; data were additionally searched against cRAP (contaminants dataset); only peptides with an ion score > 20 were considered as identified. After this second search, additional AMELY peptides were found in three individuals previously estimated as females, namely

RD9, RD14, and RD16. All the proteomic sexes agree with DNA estimation (Supplementary Table 5).

**Supplementary Table 5**. IDs correspondence and AMELY presence (●)/absence (empty cell) after MaxQuant (MQ) and Mascot searches; females only (after MQ search) were searched with Mascot; final proteomic sex estimations were performed as a combination of the two search results.

| *Sample ID* | *Proteomic ID* | *DNA sex* | *AMELY_BOVIN MQ search* | *AMELY_BOVIN Mascot search* | *Final proteomic sex estimation* |
|---|---|---|---|---|---|
| RD_8156 | RD18 | n.d. | | | F |
| RD_8158 | RD17 | F | | | F |
| RD_8159 | RD27 | M | ● | n.d. | M |
| RD_8161 | RD13 | F | | | F |
| RD_8163 | RD15 | F | | | F |
| RD_8168 | RD14 | n.d. | | ● | M |
| RD_8173 | RD11 | n.d. | ● | n.d. | M |
| RD_8175 | RD16 | n.d. | | ● | M |
| RD_8177 | RD26 | n.d. | | | F |
| RD_8179 | RD9 | M | | ● | M |
| RD_8184 | RD10 | n.d. | ● | n.d. | M |
| RD_8185 | RD12 | n.d. | ● | n.d. | M |
| RD_8189 | RD3 | n.d. | ● | n.d. | M |
| RD_8190 | RD20 | F | | | F |
| RD_8191 | RD1 | n.d. | ● | n.d. | M |
| RD_8192 | RD28 | M | ● | n.d. | M |
| RD_8193 | RD2 | n.d. | | | F |
| RD_8194 | RD29 | M | ● | n.d. | M |
| RD_8195 | RD4 | n.d. | ● | n.d. | M |
| RD_8198 | RD21 | n.d. | ● | n.d. | M |
| RD_8202 | RD7 | M | ● | n.d. | M |
| RD_8203 | RD6 | M | ● | n.d. | M |
| RD_8204 | RD5 | F | | | F |

Deamidation levels were calculated following the approach of[47], but developing an R script ad-hoc (R version 4.0.5, "Shake and Throw", doi: 10.5281/zenodo.15024003). In brief, the number of deamidated N and Q were counted from the ‘evidence.txt’ file (MaxQuant output) for all the peptide-to-spectrum matches (PSM) and normalized to the number of unmodified N-Q. These values were grouped by peptides per charge state and a weighted average was calculated based on the intensity of each PSM. Values were then averaged by peptide. The average deamidation rate and confidence interval per sample were bootstrapped (n = 1000), using the previously-obtained peptide average values. Overall, both modern and fossil samples show highly deamidated Q residues (median > 0.90; with 1 = fully deamidated, 0 = non-deamidated) and variable deamidation rates of N residues (Supplementary Figure 6). Notably, a modern sample (ibex_mod_M1) shows the lowest median N deamidation rate of the dataset (0.62); yet, the other modern samples (ibex_mod_M2 and ibex_mod_F1) show N deamidation rates akin to fossil samples. This evidence agrees with previous works, where the deamidation rate of modern samples is highly-variable and eventually showing high rates[48]. Peptides’ lengths of AMBN, AMEL, and ENAM have been estimated for fossil and modern samples, with the latter showing a higher modal peptide length (14 AAs; Supplementary Figure 7) as expected. In addition, compared to modern, fossil samples show a higher number of short peptide chains (<14 AAs), possibly due to diagenetic hydrolysis. All the raw data (including a blank), the MaxQuant evidence file, and the deamidation script were uploaded to Zenodo (https://doi.org/10.5281/zenodo.15024003).

**Supplementary Figure 6**. Asparagine (N) and glutamine (Q) deamination rates for modern (*mod*) and fossil enamel specimens of *C. ibex*, as calculated from the ‘evidence.txt’ MaxQuant output; facets are ordered based on the median N deamidation rate. Boxplots represent the bootstrapped dataset after generating n = 1000 (re)samples with replacement for each individual. The number of peptides used for calculations is given next to the sample name.

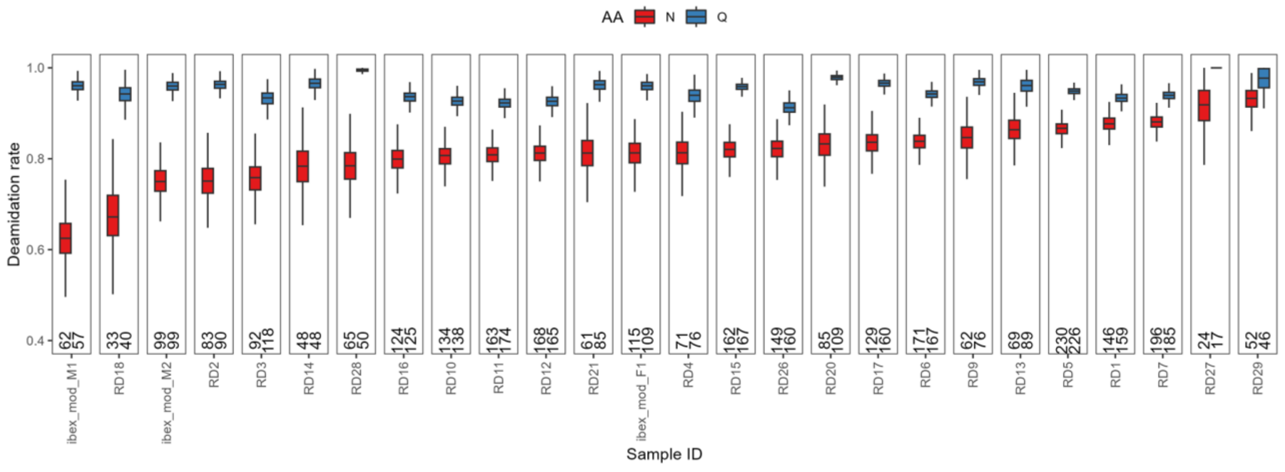

**Supplementary Figure 7.** Lengths (number of AAs) of AMBN, AMEL, and ENAM peptides for fossil and modern samples of *C. ibex*. Facets are ordered based on the modal peptide length of each sample. Red triangles represent statistical modes.

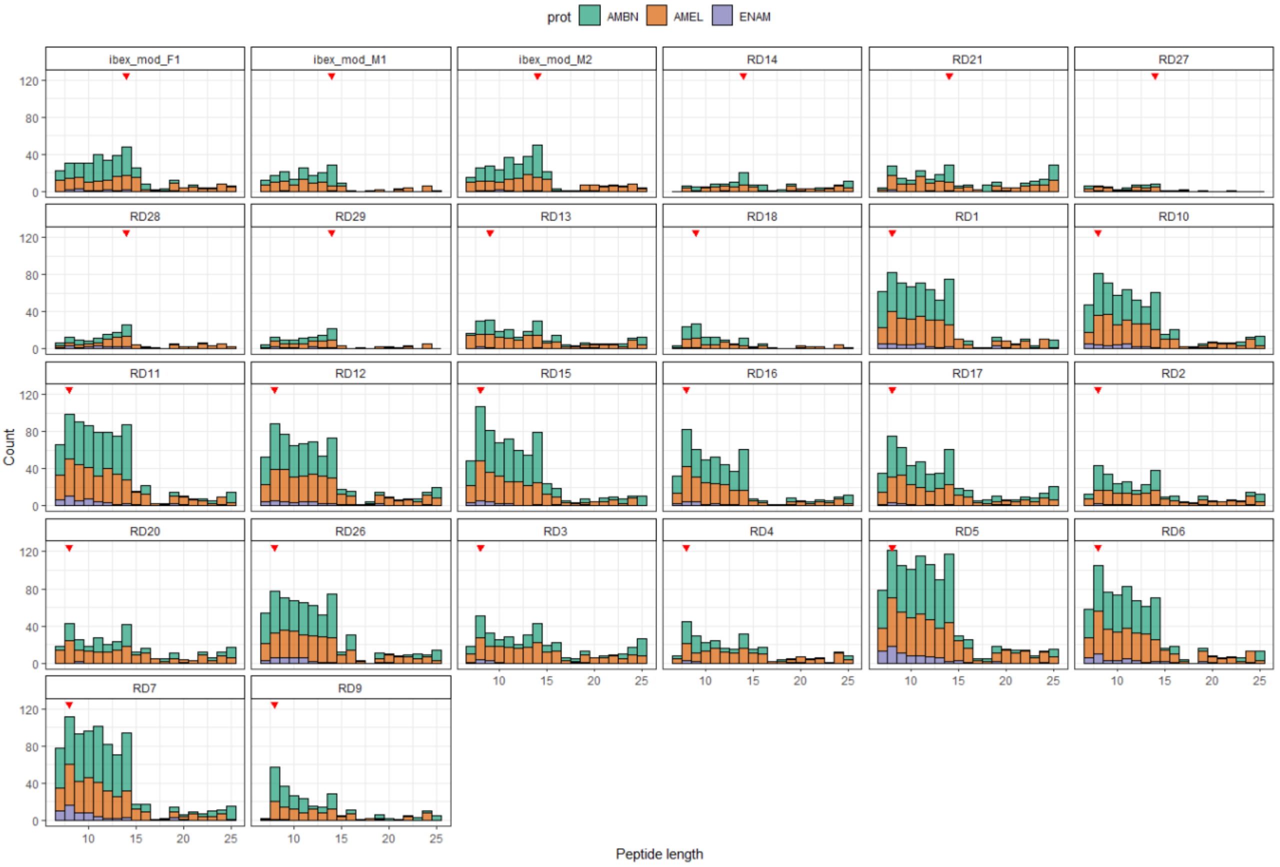


### Supplementary Note 5: Genetic sex estimation

*(Fontani, F., Iacovera, R.)*

The genetic sex of the newly generated ibex data was first estimated by applying the so-called "Mittnik method"[49]. We generated mapping statistics from reads aligned to the whole genome sequence of the *Capra hircus*, Saanen breed (GCA_015443085) and calculated the normalized ratio of sequences mapping to each chromosome. We then compared the mean ratio of reads mapping the autosomes to the X chromosome (Rx) and generated sex estimates (Supplementary Table 6) by using an edited version of the script provided in[50].

**Supplementary Table 6**. Results of Rx calculation for sex assignment

| *ID* | *Rx* | *95% CI* | *Sex assignment* |
|---|---|---|---|
| RD8158 | 0,734 | 0.687 – 0.782 | Not assigned |
| RD8159 | 0,487 | 0.477 – 0.498 | Male |
| RD8161 | 0,933 | 0.923 – 0.942 | Female |
| RD8163 | 0,841 | 0.817 - 0.865 | Female |
| RD8179 | 0,517 | 0.509 – 0.525 | Male |
| RD8190 | 0,964 | 0.956 – 0.973 | Female |
| RD8192 | 0,498 | 0.491 – 0.505 | Male |
| RD8194 | 0,499 | 0.494 - 0.504 | Male |
| RD8202 | 0,528 | 0.520 – 0.535 | Male |
| RD8203 | 0,583 | 0.537 – 0.629 | Possibly XY but not XX |
| RD8204 | 0,860 | 0.828 – 0.893 | Female |

To further validate the genetic results, we first replicated the methodology presented in[51] to calculate the Y-chromosomes to autosomes read depth ratio observed in each sample after mapping to the annotated genome of *C. hircus* Saanen. A standardized mean read depth was initially calculated by computing (read number/chromosome length)*100. Thus, we expected a ratio of standardized mean read depth of the autosomes to X chromosomes to be equal to 1 for female individuals. For males, an expected 2:1 ratio for read depth of autosomes compared to read depth on Y- and X-chromosomes was calculated (Supplementary Table S3). We then used the "Skoglund method" to calculate the ratio of reads mapping to the Y-chromosome, manually setting the Ry male limit to 0.025 (Supplementary Table 7).

**Supplementary Table 7.** Results of Ry calculation for sex assignment

| *ID* | *NchrY+ NchrX* | *NchrY* | *Ry* | *SE* | *95% CI* | *Assignment (–malelimit=0.025)* |
|---|---|---|---|---|---|---|
| RD8158 | 11751 | 313 | 0,026 | 0,001 | 0.023-0.029 | Possibly XY but not XX |
| RD8159 | 93654 | 5082 | 0,054 | 0,0007 | 0.052-0.055 | XY |
| RD8161 | 230575 | 458 | 0,002 | 0,0001 | 0.001-0.002 | XX |
| RD8163 | 29556 | 342 | 0,011 | 0,0006 | 0.01-0.012 | XX |

| | | | | | | |
|---|---|---|---|---|---|---|
| RD8179 | 35097 | 1972 | 0,056 | 0,001 | 0.053-0.058 | XY |
| RD8190 | 33498 | 79 | 0,002 | 0,0003 | 0.001-0.002 | XX |
| RD8192 | 293567 | 17080 | 0,058 | 0,0004 | 0.057-0.059 | XY |
| RD8194 | 378320 | 21465 | 0,056 | 0,0004 | 0.056-0.057 | XY |
| RD8202 | 5282 | 370 | 0,07 | 0,003 | 0.063-0.076 | XY |
| RD8203 | 3605 | 163 | 0,045 | 0,003 | 0.038-0.052 | XY |
| RD8204 | 5674 | 77 | 0,013 | 0,001 | 0.010-0.016 | Possibly XX but not XY |
